# A novel method for comparison of arterial remodeling in hypertension: quantification of arterial trees and recognition of remodeling patterns on histological sections

**DOI:** 10.1101/424119

**Authors:** Alex A. Gutsol, Paula Blanco, Svetlana I. Samokhina, Sergey A. Afanasiev, Chris R.J. Kennedy, Sergey V. Popov, Kevin D. Burns

## Abstract

Remodeling of spatially heterogeneous arterial trees is routinely quantified on tissue sections by averaging linear dimensions, with lack of comparison between different organs and models. The impact of experimental models or hypertension treatment modalities on organ-specific vascular remodeling remains undefined. A wide variety of arterial remodeling types has been demonstrated for hypertensive models, which include differences across organs. The purpose of this study was to reassess methods for measurement of arterial remodeling and to establish a morphometric algorithm for standard and comparable quantification of vascular remodeling in hypertension in different vascular beds. We performed a novel and comprehensive morphometric analysis of terminal arteries in the brain, heart, lung, liver, kidney, spleen, stomach, intestine, skin, skeletal muscle, and adrenal glands of control and Goldblatt hypertensive rats on routinely processed tissue sections. Mean dimensions were highly variable but grouping them into sequential 5 μm intervals permitted creation of reliable linear regression equations and complex profiles. Averaged arterial dimensions demonstrated seven remodeling patterns that were distinct from conventional inward-outward and hypertrophic-eutrophic definitions. Numerical modeling predicted at least twenty variants of arterial spatial conformations. Recognition of remodeling variants was not possible using averaged dimensions, their ratios, or the remodeling and growth index. To distinguish remodeling patterns, a three-dimensional modeling was established and tested. The proposed algorithm permits quantitative analysis of arterial remodeling in different organs and may be applicable for comparative studies between animal hypertensive models and in human hypertension. Arterial wall tapering is the most important factor to consider in arterial morphometry, while perfusion fixation with vessel relaxation is not necessary. Terminal arteries in organs undergo the same remodeling pattern in Goldblatt rats, except for organs with hemodynamics affected by the arterial clip. The existing remodeling nomenclature should be replaced by a numerical classification applicable to any type of arterial remodeling.

**Author summary:** Arterial hypertension effects modern nations and is characterised by systemic hypertensive angiopathy that affects all organs. Arterial remodeling is a main factor to be analyzed in animal models and human. Despite abundant data, there is a significant lack of comparative analysis on arterial remodeling. The data from the present study have established a novel methodological approach to assess and compare arterial remodeling in hypertension. We have developed an effective algorithm for morphometry of intra-organ arteries to standardize remodeling assessment and allow comparisons between different hypertensive models, organs and species. Our study opens the possibility to assess remodeling using conventional widely used histological tissue sections with no need for special perfusion-fixation. The method will elucidate the improvement and development of animal models of hypertension, and enhance the assessment of experimental therapeutic modalities.

## Introduction

The importance of understanding the pathogenesis of hypertension is undisputed, and despite recent decreases in mortality due to heart disease and stroke, the burden of disease remains high. Based on data from 2018, 33% of adults in North America have hypertension, but only 53% of those with documented hypertension have their condition controlled to target levels[1]. An important feature of hypertensive angiopathy is vascular remodeling: a complex structural and spatial modification in small arteries that is of crucial functional consequence since it alters peripheral resistance and impairs contractility[2],[3],[4],[5].

The features of arterial remodeling in hypertension have been extensively studied[3],[4],[6],[7]. To date however, there are no data demonstrating how arterial remodeling in spontaneously hypertensive rats (SHR) is distinct from hypertension due to angiotensin II or deoxycorticosterone acetate infusion, inhibition of nitric oxide synthesis, or Goldblatt’s model. The utility of animal models must be based on their resemblance to human pathology, yet commonly used measures are unable to quantify how arterial remodeling in experimental animals corresponds to arterial remodeling in humans. Thus, the well-established clinical concept of target organ damage in hypertension[8] has yet to be supported by data indicating that arterial remodeling is more extensive in the kidney or heart, for example, compared to the skin, liver, or other organs.

In the majority of studies complex three-dimensional (3D) intra-organ arterial trees have been quantified by simple measures first described more than 80 years ago[9], consisting of the wall-to-lumen ratio (WLR), and/or the mean values for a variety of arterial dimensions, e.g., external diameter (ED), internal diameter (ID), wall thickness (WTh), external perimeter (EP), internal perimeter (IP), media cross sectional area (MCSA), lumen cross sectional area (LCSA), total cross sectional area (TCSA) and internal radius (IR). These measures are highly variable and inconsistent, and do not permit comparison of remodeling between different organs, hypertension models or species. Presumably, that quantification of tapered 3D intra-organ arterial trees on tissue sections is oversimplified, and conventional averaging of dimensions in vessel morphometry led to incomparable stochastic results. Modern micro-computed tomography achieves 3D images of peripheral vessels with high resolution: ~ 2.5 μm voxel size. However, in this technique, detailed microstructure cannot be described since only the contrast-filled lumen is visualised. While this is useful for angiogenesis studies, analysis of hypertensive remodeling is limited[10].

Arterial wall remodeling patterns have been classified as hypertrophic, hypotrophic or eutrophic, associated with inward narrowing or outward widening of the lumen[6],[11]. To date, however, there is significant discrepancy with regards to the type of remodeling that develops in terminal arteries (TAs) of different organs in various hypertensive models. In SHR for example, renal arterioles may demonstrate remodeling characterized as outward hypertrophic[12],[13],[14], inward hypertrophic[15], solely hypertrophic[13] or no change[14],[16]. Some authors conclude that renal arteries > 60 μm do not develop remodeling[17], while others suggest an absence of remodeling for smaller arteries < 60 μm[16]. Mesenteric arteries may show inward hypertrophy or no change[12],[18]. Similar apparent contradictions appear in other hypertensive models[19],[20],[21],[22].

Such discrepancies could arise since arterial remodeling has previously been classified using empirical drawings, without precise quantitative analysis[11]. Since it was introduced twenty years ago, the classification has been extensively reviewed[5],[23],[24],[25] but has not been challenged with quantitative methods. Furthermore, frequently used parameters, such as WLR, remodeling index (RI), and growth index (GI) have not been rigorously tested as markers of remodeling. We therefore set out to i) study the classification of arterial remodeling patterns in hypertension, using mathematical methods, and ii) elucidate how arterial remodeling differs across a variety of organs. An algorithm for arterial remodeling assessment was developed, and we then determined if it could distinguish variants of remodeling in different organs within one model of hypertension in Goldblatt one-kidney one-clip (1K1C) rats. We hypothesized that, if conventional averaging was avoided, all organs would show the same arterial remodeling pattern. We also hypothesized distinct remodeling patterns for the adrenal gland and kidney, where the 1K1C model creates particular hemodynamic conditions. In this model, arteries within the adrenal glands experience the effects of an activated renin-angiotensin system (RAS), similar to other organs, but also experience enhanced flow due to diversion of blood from the main renal artery as a result of distal stenosis or ligation, similar to hemodynamic models of overflow[26],[27]. The remaining kidney also experiences the vasoconstrictive effects of an activated RAS, but under low blood flow due to the clipped renal artery[28],[29], that corresponds to low blood flow models[26],[30].

We also determined if examination of random tissue sections can provide useful information in studying hypertension, compared to use of arterial myography[3],[31]. In this regard, the vast majority of data from *in vitro* myography experiments has been generated from dissected mesenteric arteries, and may not be applicable to other organs. Finally, studies have recommended that arterial morphometry should only be performed on perfusion-fixed organs with pharmacologically relaxed vessels[31],[32]. We therefore examined if routine immersion fixation is appropriate for arterial remodeling analysis to elucidate comparisons between animal and human samples, since for the latter immediate relaxation and perfusion fixation are not practical in general.

## Results and Discussion

### Part I. Quantification of arterial trees on histological sections

#### Means of linear sizes and ratios are not applicable for arterial morphometry on tissue sections

While the parenchyma of different organs is well represented on histological sections, arteries are an exception. They appear as circles and irregular strips with a wide range of sizes and shapes, which are difficult to quantify[33]. To date, there are two approaches to arterial morphometry. *In vitro* myography uses similar vascular segments to minimize variability and allow comparison of averaged dimensions or their ratios among different models, or different arteries in the same model[34],[35]. However, myography data are restricted to only a few areas that are suitable for sampling - mainly the mesentery and aorta. Second, quantification of intraorgan arteries is based on casual measurements of arterial segments on tissue sections. However, the majority of studies simply average the sizes and ratios, as for myography data, resulting in a high degree of variability and difficulty in comparing organs. Measured vessels are defined with uncertain ranges. For example, for the kidney: ‘arteries and arterioles’[36]; ‘intrarenal’[37],[38]; ‘proximal interlobular’[13],[39]; ‘afferent arterioles’[40],[41]; ‘vessels adjacent to glomeruli in the outer cortex’[42] or ‘at the same level proximal one-third from corticomedullary junction’[43]. Similar ‘interlobular arteries’ could vary significantly in ID ( ~100 μm[44], ~40 μm[13], ~25-50 μm[45], ~30-250 μm[22]).

TAs from multiple organs in normal rats were first analyzed using that conventional approach with measurements of mean ED, ID, WTh and WLR (**Table 1**). All values were obtained within an arterial ED range of 10-50 μm, since larger or smaller arteries were difficult to find in sufficient numbers in tissue samples. TAs in each organ were characterized by distinct average linear dimensions and WLRs. For instance, although bronchial and skeletal muscle arteries demonstrated similar WLRs, their EDs, IDs and WThs differed (P < 0.01). The WLRs for brain arteries were lower but IDs larger than in liver arteries (P<0.001), although EDs and WThs were similar between these two organs. Importantly, coefficient of variation of arterial dimensions were relatively high, varying between 40-70%, indicating their significant diversity[46]. Accordingly, individual histograms for parameter distribution in each organ were prepared. Histograms for ED, ID, and WTh demonstrated considerable variability, without a normal Gaussian distribution (**Fig 1**). Indeed, as shown in **S1 Table**, the extent of variability for ED, ID and WLR measures did not pass conventional statistical tests for normality.

**Table 1.** Averaged linear dimensions and wall-to-lumen ratio for terminal arteries with ED of 10 – 50 μm in different organs.

| **Organ** | **ED,**<br>**$\mu\text{m}$** | **ID,**<br>**$\mu\text{m}$** | **WTh,**<br>**$\mu\text{m}$** | **WLR,**<br>**%** |
| --- | --- | --- | --- | --- |
| **Liver** (N=63) | 21.6±1.4 | 8.8±0.8** | 6.4±0.4 | 98.6±7.2** |
| CV | 54% | 69% | 48% | 58% |
| **Adrenal** (N=61) | 22.0±1.1 | 8.7±0.5 | 6.6±0.4 | 81.9±5.1 |
| CV | 40% | 41% | 52% | 48% |
| **Kidney** (N=148) | 32.7±0.6 | 13.3±0.4 | 9.7±0.2 | 78.9±2.2 |
| CV | 24% | 33% | 24% | 35% |
| **Skin** (N=225) | 20.1±0.6 | 8.5±0.3 | 5.8±0.2 | 75.3±1.9 |
| CV | 48% | 52% | 52% | 37% |
| **Skeletal muscle** (N=98) | 19.7±0.7* | 8.4±0.3* | 5.6±0.2* | 70.8±3.1 |
| CV | 37% | 37% | 45% | 43% |
| **Bronchial arteries** (N=50) | 37.1±1.3 | 18.4±1.2 | 9.4±0.4 | 68.7±7.5 |
| CV | 24% | 48% | 32% | 77% |
| **Heart** (N=115) | 26.6±0.9 | 12.7±0.6 | 6.9±0.3 | 63.6±3.0 |
| CV | 40% | 52% | 42% | 50% |
| **Spleen** (N=203) | 17.7±0.5 | 8.3±0.3 | 4.7±0.1 | 61.6±1.3 |
| CV | 42% | 51% | 38% | 30% |
| **Small Intestine** (N=55) | 18.9±1. | 9.5±0.8 | 4.7±0.2 | 57.2±3.0 |
| CV | 42% | 65% | 38% | 39% |
| **Stomach** (N=90) | 24.9±1.0 | 12.7±0.6 | 6.1±0.3 | 53.5±3.0 |
| CV | 41% | 46% | 54% | 54% |
| **Brain** (N=144) | 23.4±0.6 | 12.7±0.4 | 5.4±0.2 | 45.6±1.7 |
| CV | 32% | 40% | 35% | 46% |
| **Pulmonary arteries** (N=66) | 29.2±1.3 | 20.9±1.0 | 4.1±0.2 | 21.9±1.1 |
| CV | 35% | 41% | 34% | 42% |

Data are mean ± SEM. CV - coefficient of variation; N - number of measured arteries; \*P<0.01 vs the bronchial arteries; \*\*P<0.001 vs the brain. Organs are displayed in order of descending WLR values.

**Fig 1.**
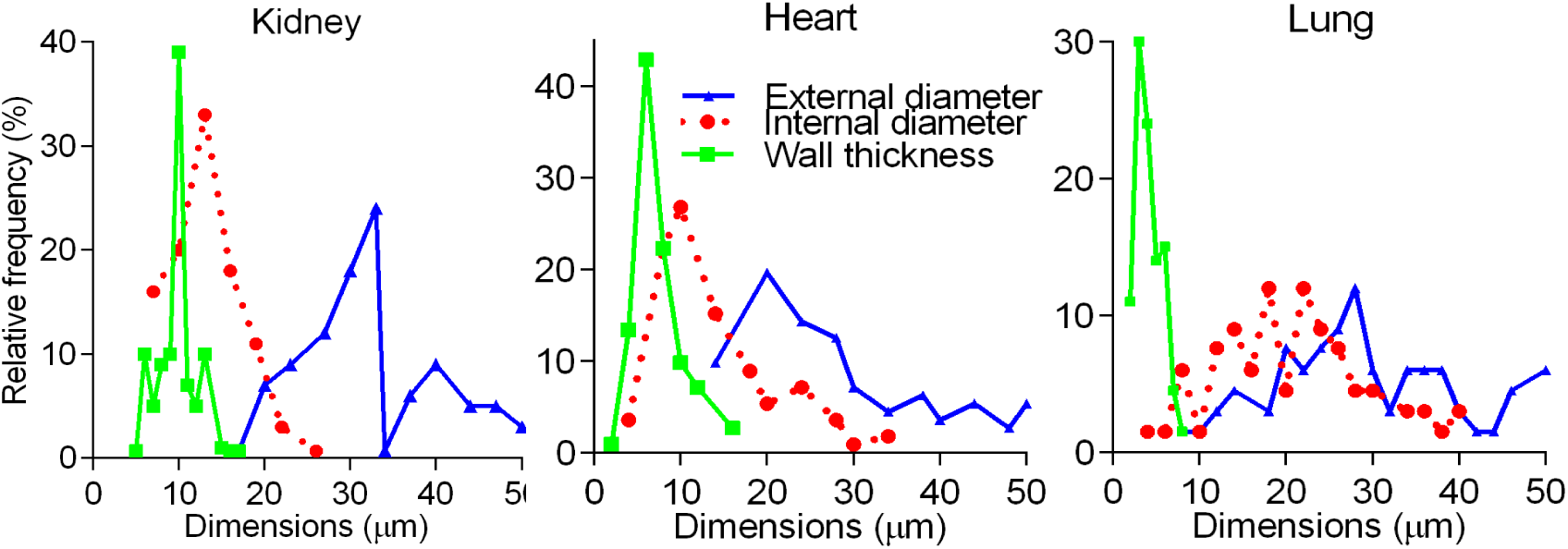
Detailed statistical analysis of primary data. Histograms for ED, ID, and WTh in the kidney, heart and pulmonary arteries, used to calculate statistics in Table 1, demonstrated irregular frequencies and profound asymmetry.

Our detailed analysis (**Table 1, Fig 1, S1 Table**) indicates that mean values have little statistical value with regards to such measurements[46], since even a small range of EDs (10-50μm) for TAs in all organs was associated with an abnormal distribution and high variability. Furthermore, increasing numbers of measurements to normalize distribution and reduce variability are not effective for tapering branching objects[33].

While WLR is considered independent of diameters and not affected by the size of arteries studied[3],[9],[41], this matter has not been rigorously substantiated in the literature[4],[5],[25],[32],[47]. Furthermore, we found as many as sixteen different formulae to calculate WLR. Some studies employed the diameter or radius: WTh/ID[20],[44],[48]; ED - ID/2ID[49]; ED/ID[50]; WTh/IR[12][17]; a media index WTh/IR with IR specifically defined as the distance from the center of the arterial lumen to the middle point of the media[51]; 2WTh/ED[52],[53]; 2WTh/ID[54]; WTh/ED[55]; or ID/ED[56]. Others have measured WLR from the perimeters as EP/IP[40],[41],[42], or preferred to use the area: MCSA/LCSA[15],[16],[45]; MCSA/TCSA[37]; LC SA/MC SA[36]; LCSA/TCSA[57]. The WLR has also been interpreted as the ratio of the area to perimeter MCSA/IP[22], or renamed RI[58]. Rarely, point counting has been used to estimate WLR as the average ratio of the volume density of the walls to the volume density of the lumens[16],[59]. However, many studies indicate that WLR is not constant but declines 2-10-fold in arteries of ED 20-500 μm in the human and rat kidney[51],[60]; human and rat brain[61],[62]; and human and dog liver[48],[63]. Our data indicate that WLR is not constant even within a small range of EDs (**Fig 2A**).

**Fig 2.**
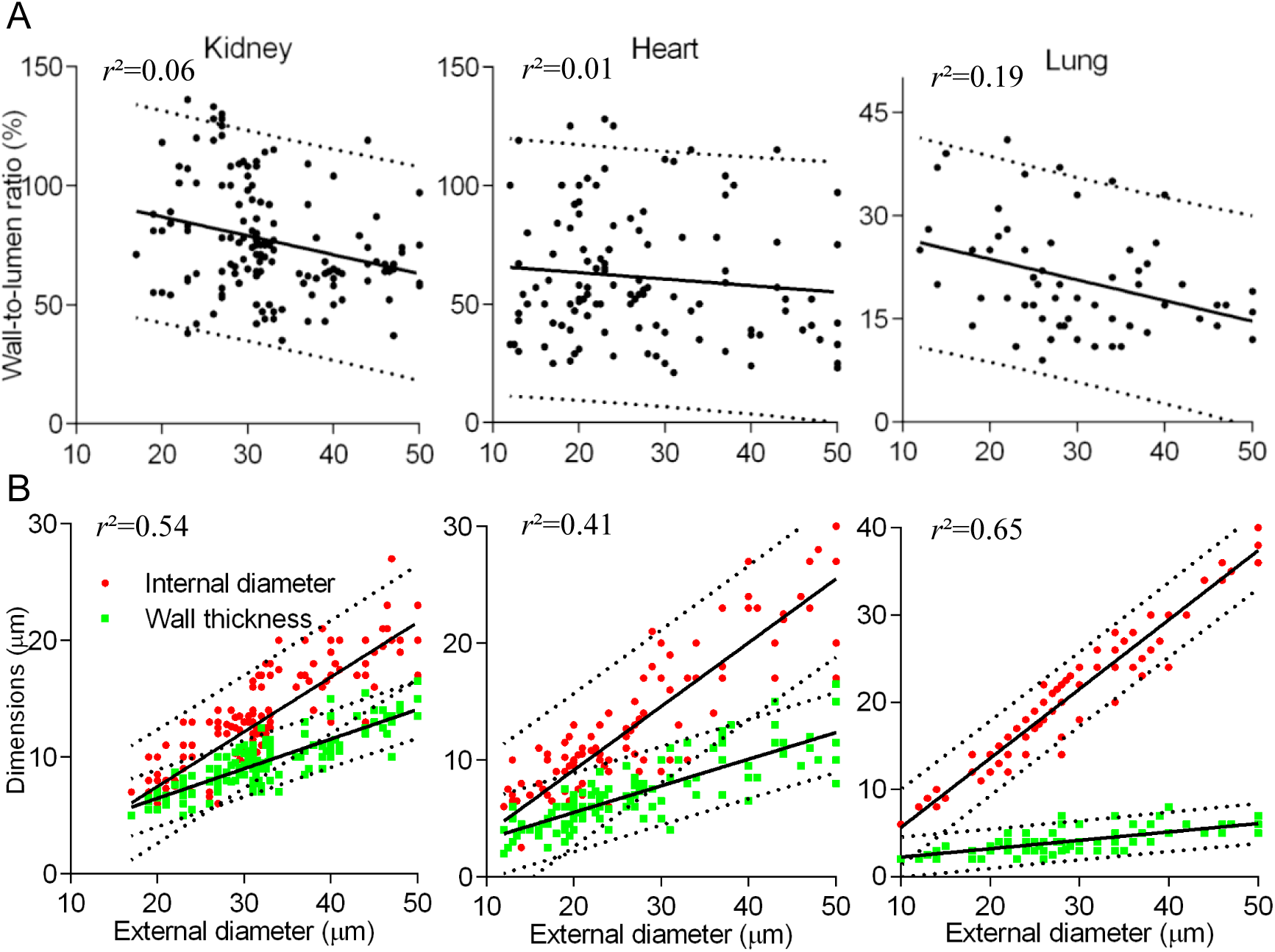
Detailed statistical analysis of primary data. (A) WLR scatterplots were irregular with low *r*^2^ (solid lines), and wide 95% prediction bands (dashed lines). Corresponding statistics are shown in **Table 1**. (B) Scatterplots of primary data displayed certain incremental change in arterial dimensions. Coefficients *r*_2_ were moderate, and the pulmonary arteries had the best fitting value.

In summary, neither averaged WLR nor the mean values for ED, ID or WTh are applicable to even relatively small intervals of vessel caliber. Thus, any comparison of mean values for these measures is not appropriate[46].

#### Application of short intervals provides precise complex profiles and linear regression equations

For vessels with EDs in the range of 10-50 μm, there was a trend towards dependence of ID and WTh on ED, although variability was substantial (**Fig 2B**). We therefore assessed primary data with a complex profile method, which is widely used in cartography, thermodynamics, and engineering[64],[65]. In this technique, a contour line is drawn as a function of two variables, where the gradient of function is perpendicular to the contour isoline, thus representing more than two dimensions on a two-dimensional (2D) graph. To be applied to measuring arteries, each arterial circle on a histological section is first characterized by three values: ED, ID, and WTh. Then all measured vascular circles were arranged in a row in order of ascending ED, and numbered from 1 to N, where N is the total number of measured arterial rings. Subsequently, each number was plotted on the y-axis, while its corresponding ED, ID and WTh were plotted on a bidirectional x-axis (**Fig 3**).

**Fig 3.**
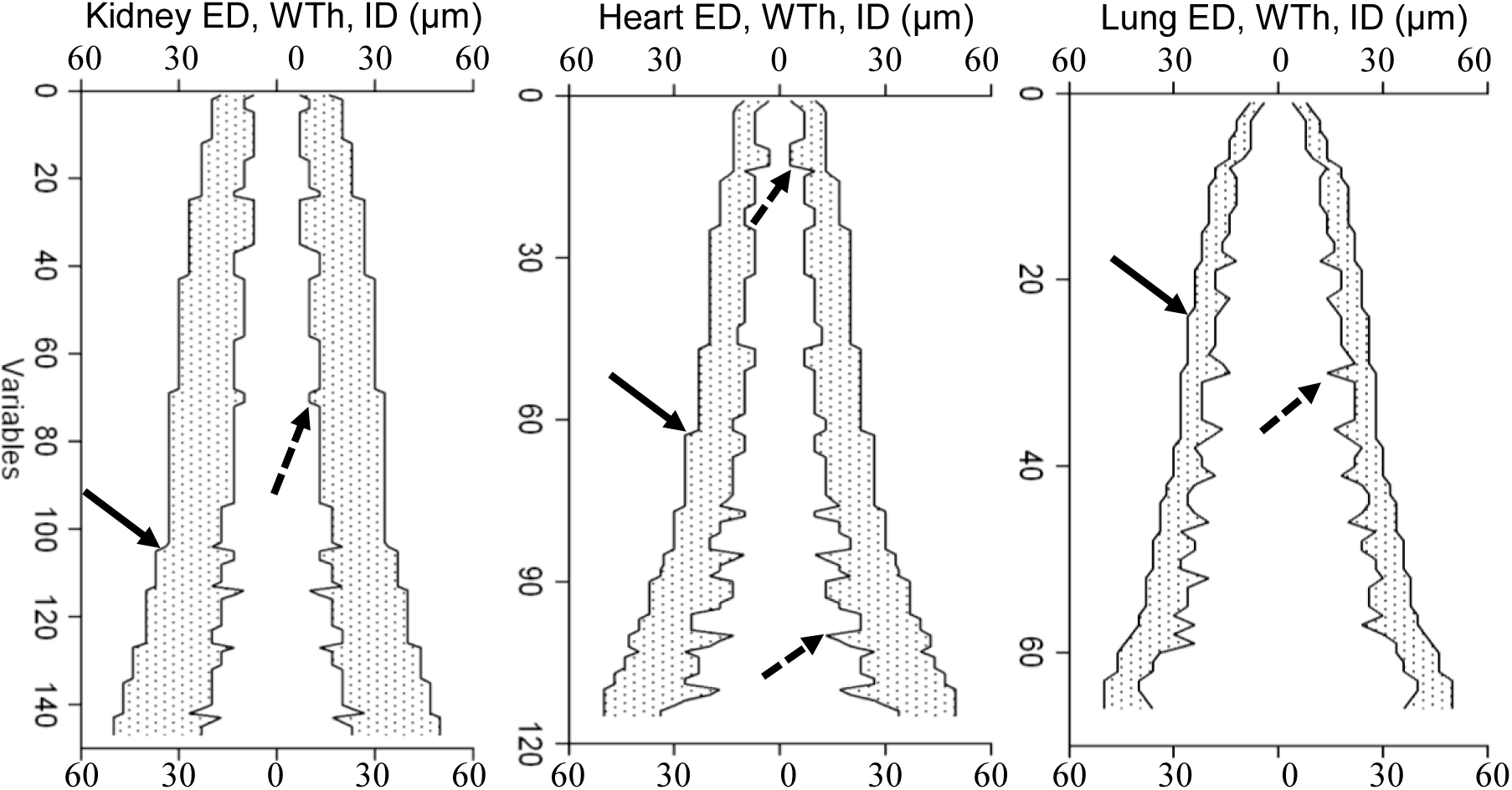
Analysis of primary data with complex profiles. Primary measurement for the kidney, heart, and pulmonary arteries were arranged in the complex profiles. Axis X – the bidirectional common scale for ED (outer contours), ID (inner contours), WTh (shaded regions); axis Y represents the number of measured arteries (variables) in order of ascending ED. Steps in the outer contours (solid arrows) reflected a minimal division on an eyepiece micrometer. Outliers along the internal contours (dashed arrows) corresponded to wider or narrower ID, compared to neighboring ED values.

The same primary data, represented in this way, revealed a regular hemodynamic structure. Staircase steps of the external contours were relatively small and regular since a conventional microscope with a 40x objective provided a minimal division value of ~2 μm to the eyepiece micrometer, so that each ED was rounded off to ± 2 μm. Spikes of internal contours were larger and quite irregular corresponding to some arterial segments with similar ED but different ID and/or WTh, and it was necessary to understand the source of multiple outliers. Presumably, those outliers represented subsets among arterial segments, branching at different hydrodynamic points. We speculated that arteries with similar ED at proximal regions should have thicker media, i.e. higher wall-to-lumen ratio, in response to larger pressure gradients (**Fig 4A**).

**Fig 4.**
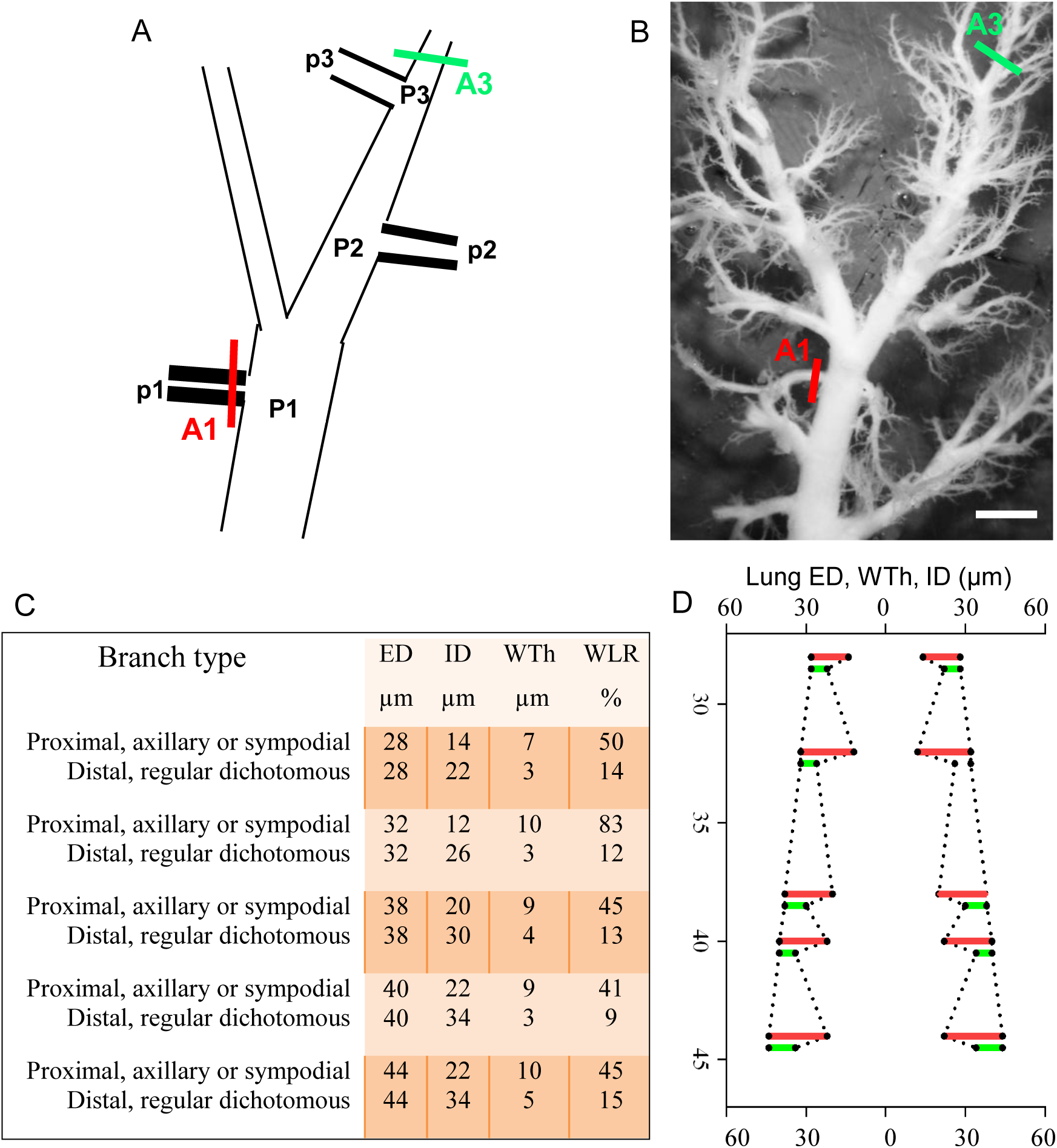
The hemodynamic explanation for the appearance of outliers. (A) According to Fick’s law, the capillary pressure in proximal (p1) or distal (p2, p3) regions must be similar, while arterial pressure in a proximal artery P1 is higher than in distal arteries P2 and P3. Consequently, Δ(P1- p1) > Δ(P3 - p3). Arteries at proximal regions should have thicker media, i.e. higher wall-to-lumen ratio, in response to larger pressure gradients. (B) The paired sampling of pulmonary arteries. Forty five arterial pairs with the same ED were identified in the proximal axillary and distal dichotomous branches respectively. For example, A1 (the red bar) and A3 (the green bar), sampled into paraffin blocks, and dimensions were measured on stained sections. Scale bar, 100 μm. (C) Five pairs in the table are the examples of the total (N=45) measured pairs of branches with the same ED that could have differences from 50-110% in ID, 100-230% in WTh, and 250-600% in WLR (P<0.001). (D) In the complex profile, those neighboring branches create most outliers, as in **Fig 3**.

To test this hypothesis, dissected pulmonary arteries were measured by the paired sampling: branches of the same ED were identified in the proximal and distal segments, cut at branching points, and their dimensions were measured (**Fig 4B-D**). To compare only two vessels 5 mm apart would require enormously laborious reconstruction of approximately 2000 serial sections while the method of paired branches morphometry was much more effective. Axillary and regular dichotomous branches with the same ED but smaller ID and thicker WTh would be placed closer in profiles, determined as outliers (**Fig 3**). That explained the presence of numerous outliers as an intrinsic property of arterial branching to respond to variable local hemodynamics, that complies with a conception of constrained random branching for smaller vessels [66]. Outliers are not evident in primary data plots (**Fig 2**) to be removed. Moreover, a conventionally recommended increase in number of measurements[46] simply increases the number of outliers. To make outliers more evident, instead of conventional averaging of the data for the entire range of 10-50 μm, the ED was divided into short regular intervals (**Fig 5**).

**Fig 5.**
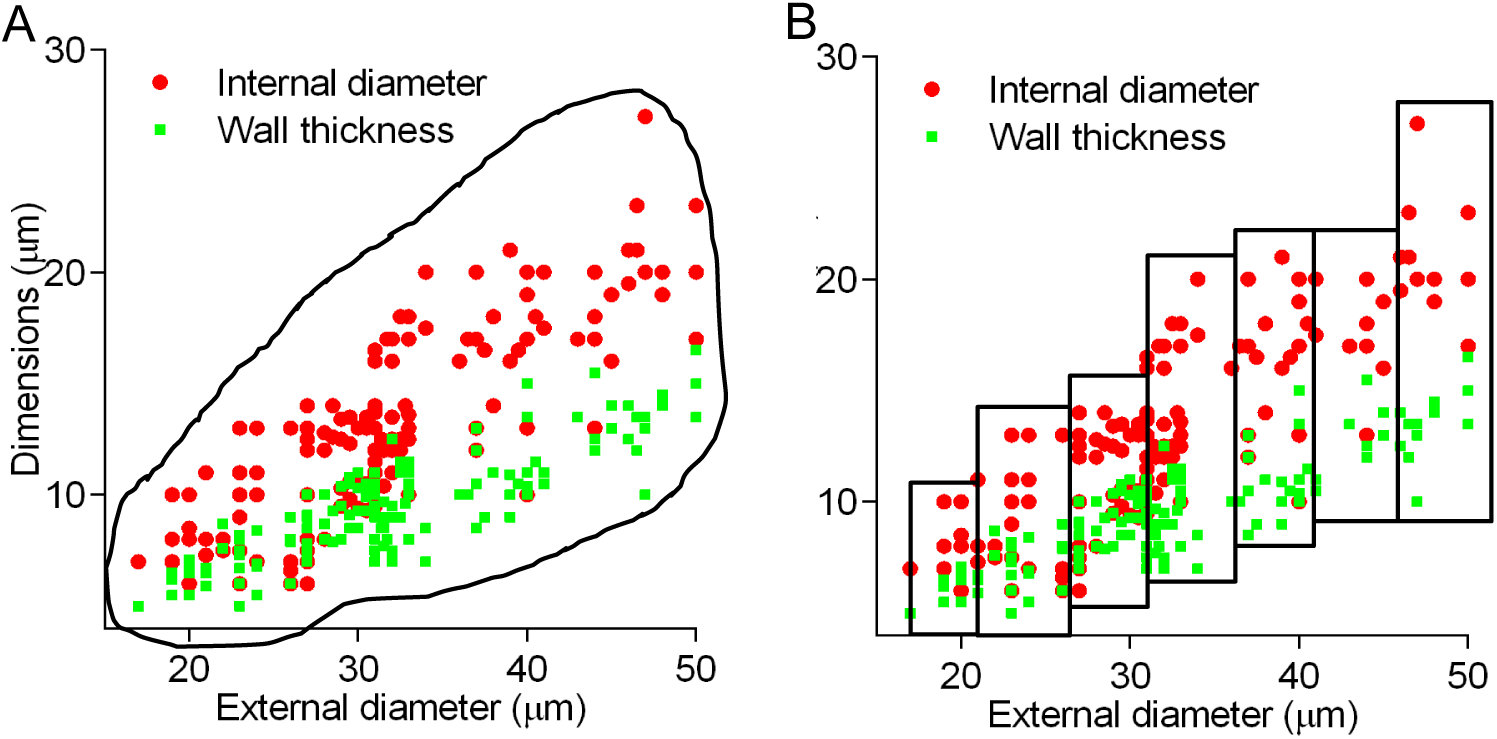
Basic approach to arterial morphometry analysis. (A) In the conventional approach, statistics of arterial dimensions are calculated for a broad range of calibers. (B) In the proposed algorithm, statistics are calculated for fixed 5μm intervals for TAs with ED ~ 10-50 μm.

To avoid averaging, some investigators have measured interval means. However, significant discrepancies in choice of interval steps and caliber ranges render such data as highly variable and difficult for comparative analysis. Some studies considered two intervals as ‘interlobular and arcuate’ with ID ~ 70 and ~160 μm[59], or ‘small and intermediate’ as ED~ 50-100 and 100-500 μm[60]. Others found it more appropriate to divide ‘interlobar vessels’ into six intervals of <70, 70-95, 95-120,and >120 μm[16]. Another variant was to cut the range of 90-220 μm into four equal classes[21]. A choice for cerebral arteries was four intervals of IR <20, 20-49, 50-79, and >80 μm but for older animals it was only 2 of 80-119 and 120-159 μm[61]. A different subdivision was used for the liver: ≤ 20, 21-50, 51-124, and > 125 μm[48]. In the lung seven non-regular intervals of 0-50, 50-100, 100-200, 500-1000, and >1000 μm were counted[53]. Thus the advantage of using intervals could be compromised by subjective choices of interval size that neither improves standardization nor comparison.

We tested different intervals, and found that the interval of 5 μm was optimal for all organs. That interval corresponds to adding one layer of vascular smooth muscle cells, which has a normal thickness of 5-7 μm[63],[67]. Indeed, the relation between WTh and the number of VSMC-layers can be represented by a linear regression line (r = 0.88)[63].

When the complex profiles had been drawn not from every measured value, but from the mean interval values, they demonstrated organ-specific gradients in tapering (**Fig 6A, S1 Fig**) allowing comparison of arterial remodeling between organs. Furthermore, in contrast to data depicted in **Fig 2B**, if outliers had been removed, the means of short intervals revealed very tight linear regression between both lumen and wall thickness and vessel caliber in kidney, heart and pulmonary vessels, and in other organs as well (**Fig 6B, S2 Fig**). All acquired equations demonstrated goodness of fit coefficients *r*^2^ between 0.8 - 0.9 (P <0.0001), with few exceptions for WTh in the brain and bronchial arteries (**S2 Table**).

**Fig 6.**
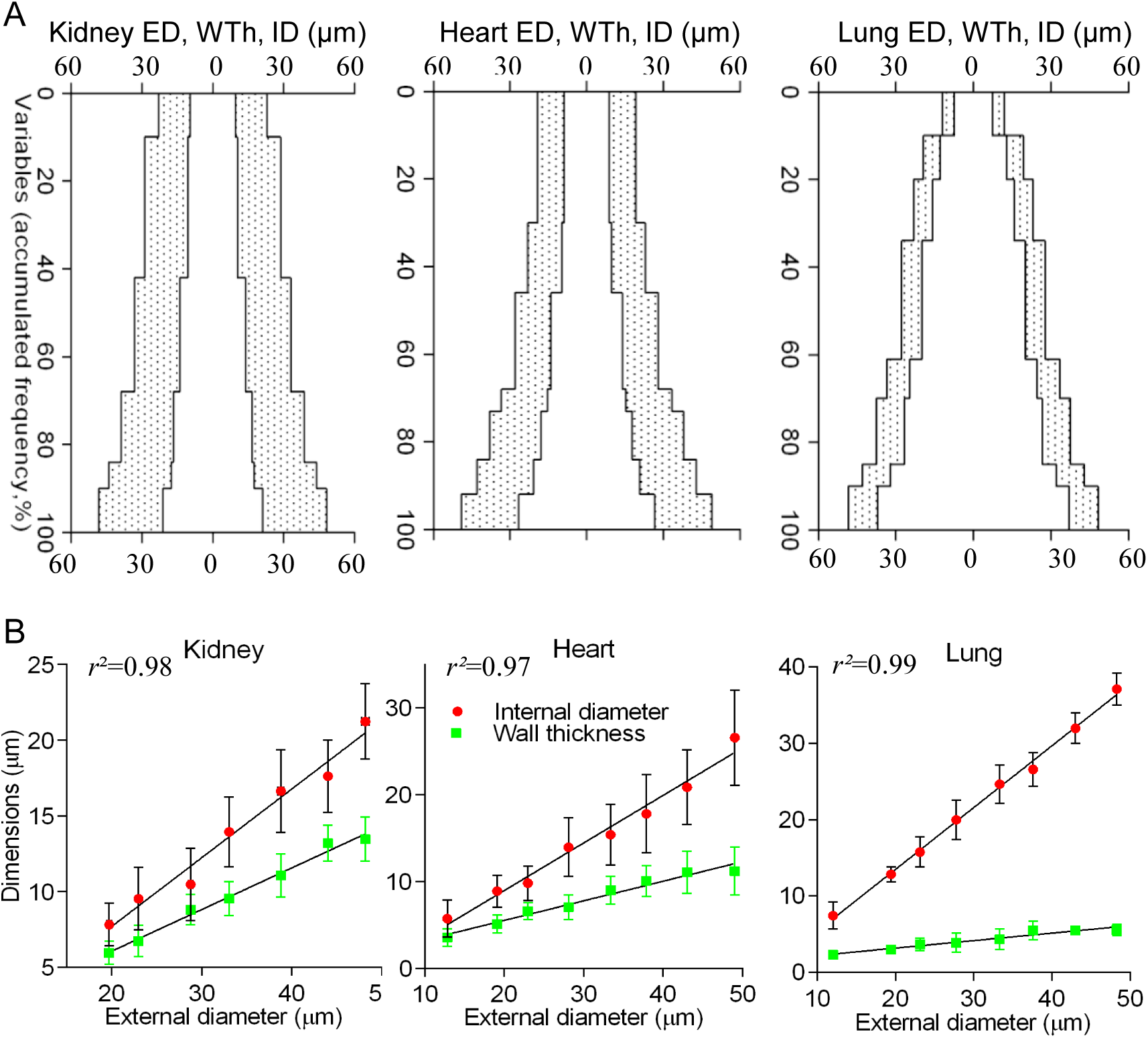
Application of the short fixed intervals significantly improved primary data analysis. (A) Complex profiles were built from the accumulated frequencies (axis Y) of the means of ED, ID, and WTh for 5-μm ED intervals. In contrast to scatterplots in **Fig 2B**, terminal arteries exhibited distinctive tapering patterns in organs. Complementary graphs are in **S1 Fig.** (B) Mean values of ID and WTh for the regular 5-μm ED intervals revealed robust linear regression equations (P<0.001). Complementary graphs are in **S2 Fig.** Points on the lines are mean ± SD for 5-μm ED intervals.

To decide whether linear regression equations for TAs were organ-specific, their slopes and intercepts were compared. Equations for pulmonary, bronchial, adrenal, stomach and skeletal muscle TAs were distinct (P<0.0001). Heart and spleen TAs had similar equations, as did brain and intestine, and kidney, liver, and skin (**S3 Fig**).

Only a few attempts to apply linear regression to arterial morphometry have been published[18],[22],[51],[63]. Those studies demonstrated that the range of EDs from 10-100 μm fits linear regression, while wider intervals of ED 100-1000 μm follow an exponential function. We found the range of EDs from 10-50 μm was the most practical to analyze. That caliber is not only the most frequently observed, but also best addresses the functional consequence of remodeling, being responsible for maximum values in peripheral resistance[3],[4],[5].

The same calculation, based on means of short intervals, was applied to compute WLR, although the ratio continued to demonstrate high variability (**Fig 7A**), presumably due to the interaction of standard deviation in the ratio of two dependent values[46]. It is evident, comparing different organs, that irregular graphs of mean interval WLRs are difficult to approximate correctly (**Fig 7B**). Therefore we calculated interval WLR from verified linear regression equations (**S2 Table**). The results demonstrated clearly that WLR is not constant, with evidence for nonlinear increases or decreases (**Fig 7C**).

**Fig 7.**
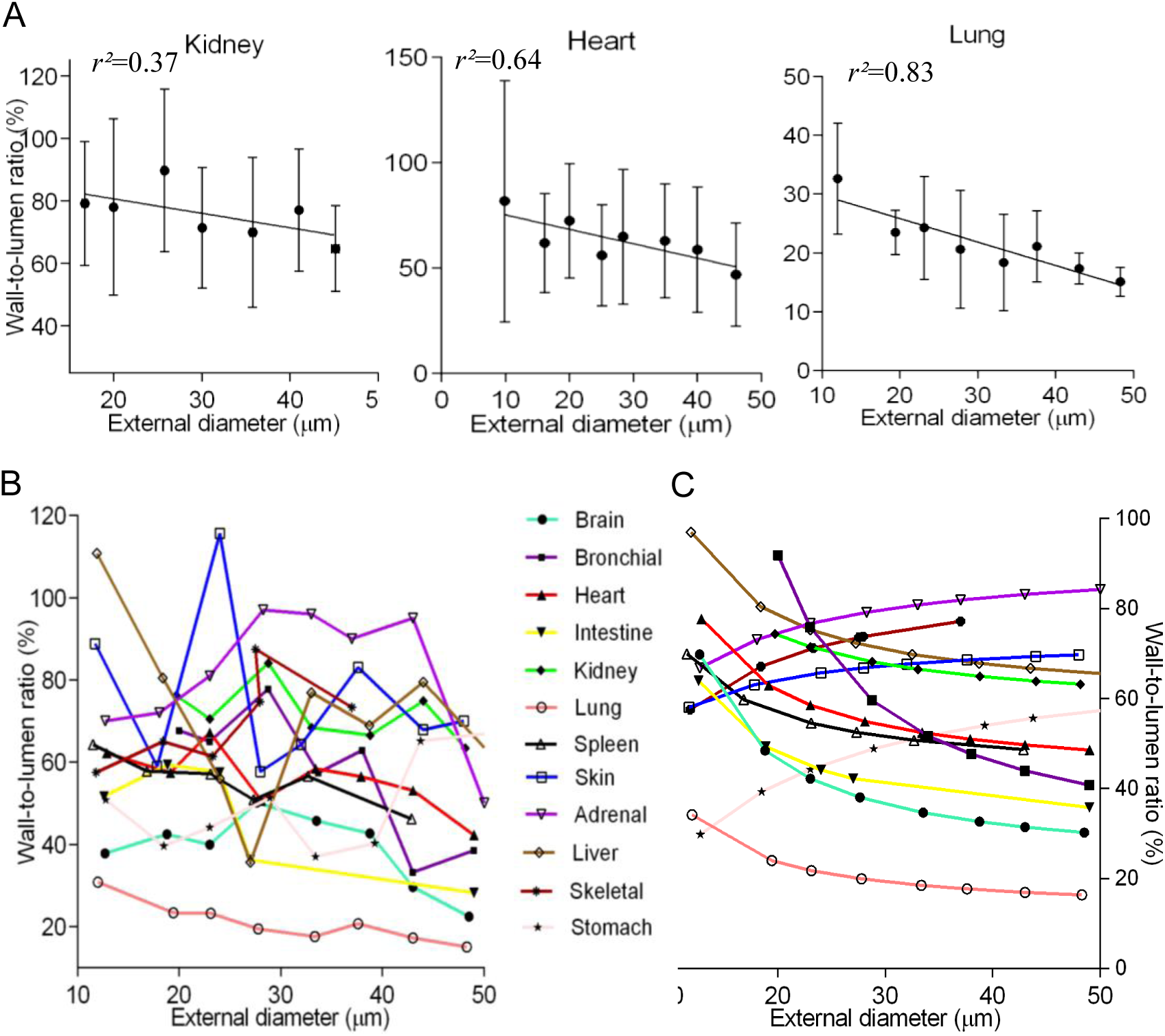
Significant variability of the wall-to-lumen ratio in different organs. (A) Interval means of WLR still varied significantly, although *r*^2^ improved as compared with Figure 2A (P ≈0.14-0.02). The pulmonary arteries had the highest *r*_2_ fitting value (P=0.002). Points on the lines are mean ± SD for 5-μm ED intervals. (B) Means of WLR for the 5-μm ED intervals were irregular, except in pulmonary arteries. (C) WLRs were counted from the linear regression equations. The ratios were not constant, decreasing or increasing by nonlinear functions.

#### Hemodynamic significance of morphometry data

Reliable linear regressions also validated an assessment of relative resistances (RR) that was calculated using Poiseuille’s relationship RR=1/πr^4^, assuming constant viscosity and length, and where r is the radius of the vessel lumen, as it was recommended[63],[68],[69]. RRs for 5 μm regular intervals across multiple organs are depicted in **Fig 8 and S2 Fig**, with the highest values in liver, and lowest in pulmonary arteries, rising exponentially in the smaller segments, since blood flow is proportional to the fourth power of the luminal radius.

**Fig 8.**
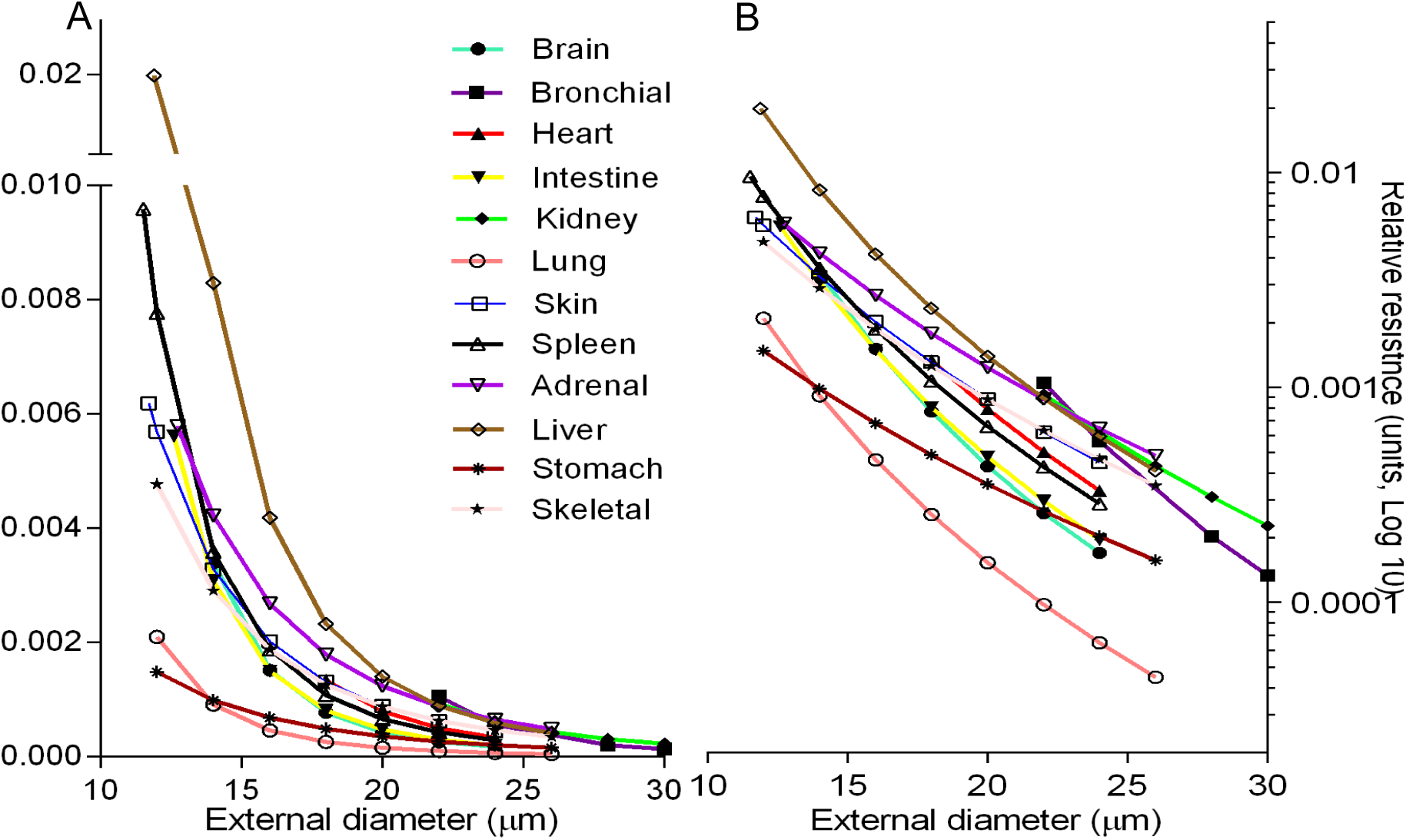
Comparative analysis of the relative resistance in different organs. (A) RR curves were calculated from the regression equations in **S2 Table.** The liver and spleen had the largest values (P<0.001), while the pulmonary and stomach arteries - the lowest (P<0.001). (B) Diversions in RRs were better revealed on the logarithmic scale.

Accordingly, terminal relative resistance (TRR) would estimate the resistance for the smallest vessels of ED= 10-20 μm that have the largest accumulated frequency in complex profiles (**Fig 9A**).

**Fig 9.**
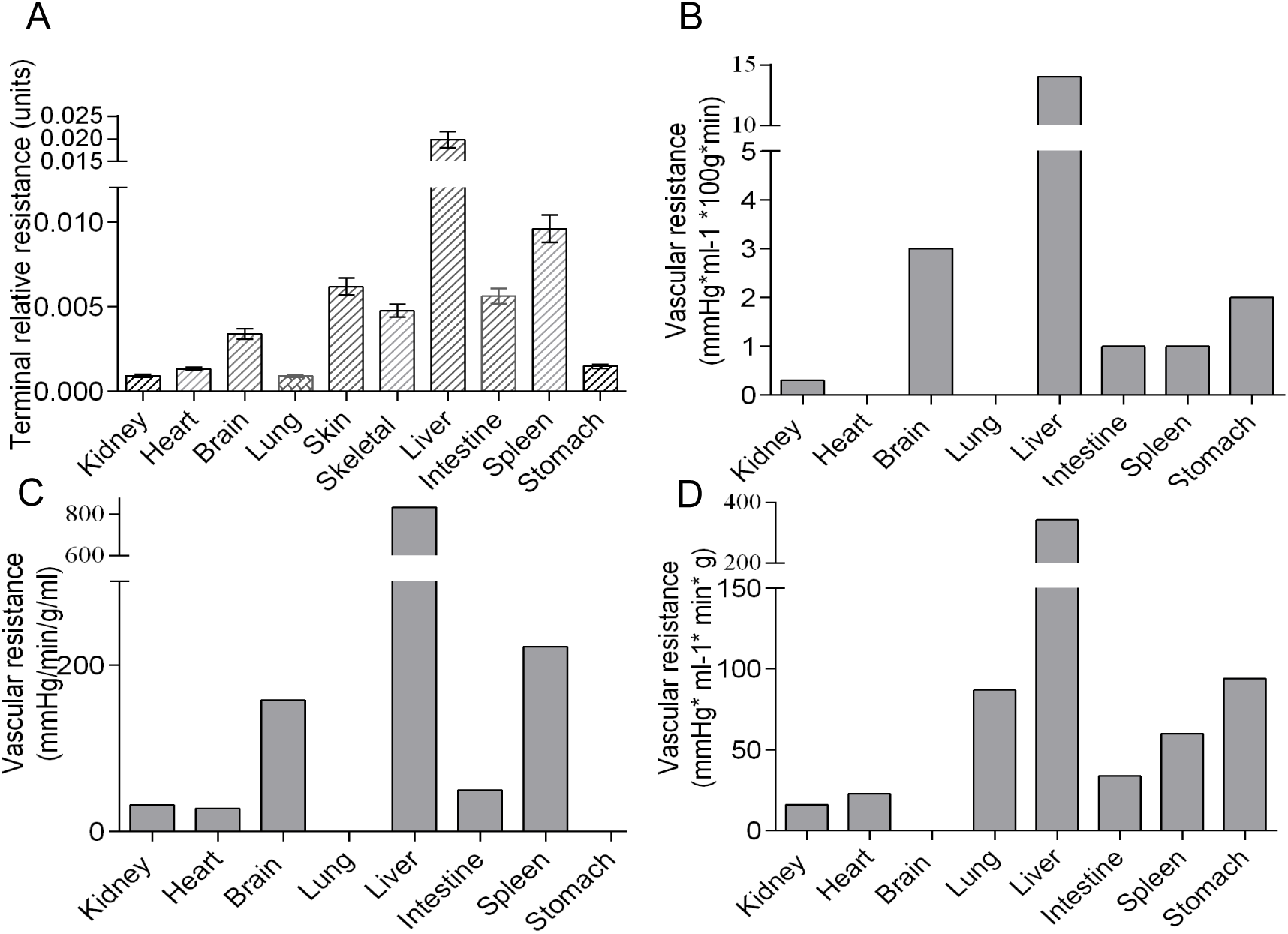
Correspondence between morphometric and hemodynamic parameters. (A) TRR demonstrated similarity in organ-to-organ ratio with vascular resistances acquired with physiological methods: modified from[70] (B); [71](C);[72] (D). Data on (A) are mean ±SEM, on (B-D) – mean values.

Since the complex profiles reveal the average numbers of arterial branches of defined lumen diameters in each organ, prior to the capillary bed, it would be reasonable to measure terminal capacity (TC) in each organ, as the sum of the internal volume of each segment multiplied by its frequency:

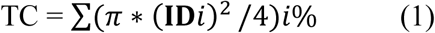

Where **ID***i* is the mean ID of each interval, calculated from the linear regression equation; *i*% - the frequency of that interval in the complex profile (**Fig 10A**).

**Fig 10.**
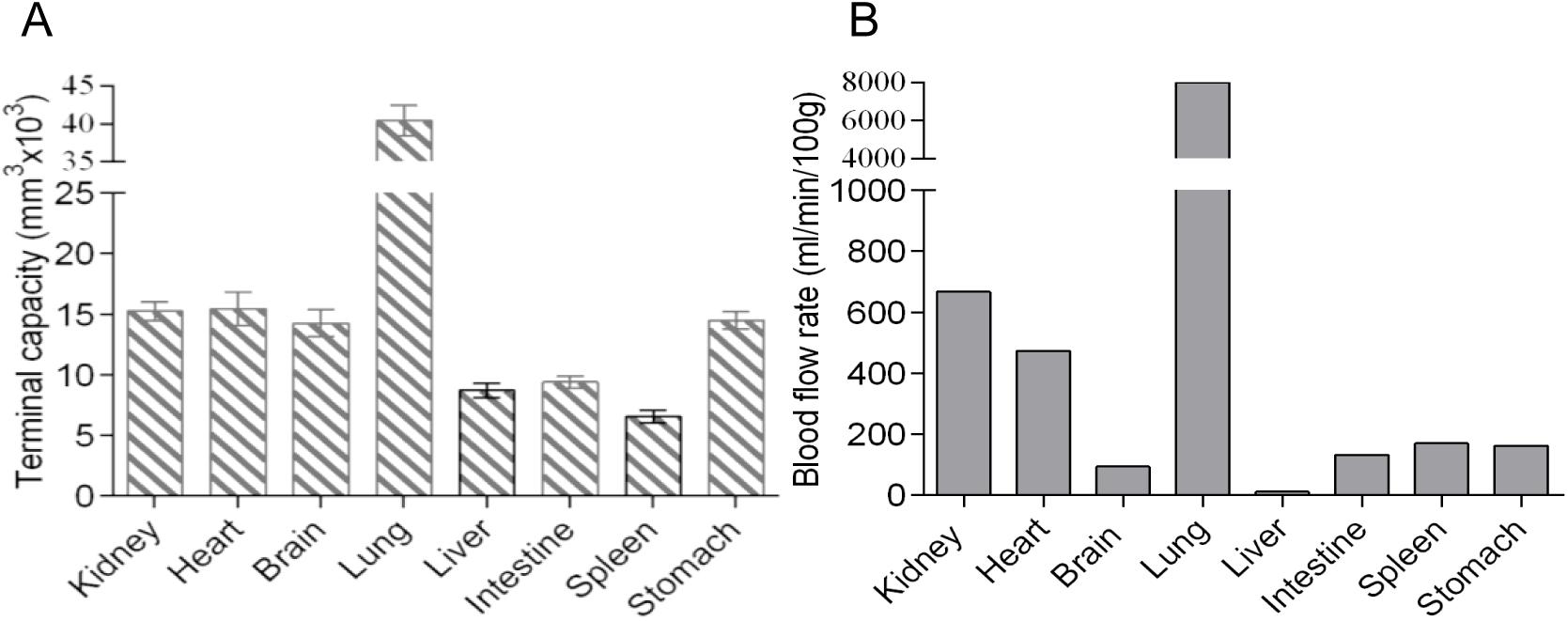
Correspondence between morphometric and hemodynamic parameters. (A) TC values were in good agreement with organ blood flow rates, estimated by a physiological method[73] (B). Data on (A) are mean ±SEM, on (B) - mean values.

The hemodynamics of each organ are characterized by blood flow rate (BFR), regional vascular resistance (RVR), vascular volume, and percentage of cardiac output received[70],[71], reflecting the uniqueness of a vascular bed. Although each organ possesses a distinctive arterial tree, there are currently no standardized morphometric parameters to quantify those distinctions and correlate them to BFR or RVR[68],[69]. Indeed averaged EDs, IDs, and WLRs are only loosely associated with hemodynamic parameters, although they accompany impaired contraction or relaxation in *in vitro* myography experiments[62]. Our algorithm provides a promising solution to this problem. The use of linear regression equations allows for quantification of TRR, which reveals significant similarity to RVRs obtained by physiological methods (**Fig 9B-D**)[70],[71],[72]. The proposed TC formula also shows good correlation with BFR, determined by microsphere method[73] (**Fig 10**).

#### Larger arteries are the most important source of errors

In TAs with ED > 40 μm, only a small number of measurements per interval could be obtained that has a crucial impact on the precision of regression equations, since even one outlier can alter the result significantly, i.e. the means of such intervals are unreliable[46], and therefore the same data set could be approximated with several equations and RR curves, depending on numbers of counted larger vessels (**Fig 11**). Small deviation of regression lines significantly affect RR according to the formula R= 1 /πr^4^. In our study, minor deviations of 8-12% in the largest IDs caused deviations of 25-90% in the smallest IDs, which in turn, had pronounced effects on the calculated TRR values, since blood flow is proportional to the fourth power of the luminal radius. To avoid such errors, the equations must be verified by comparison of corresponding RR curves (**Fig 11**).

**Fig 11.**
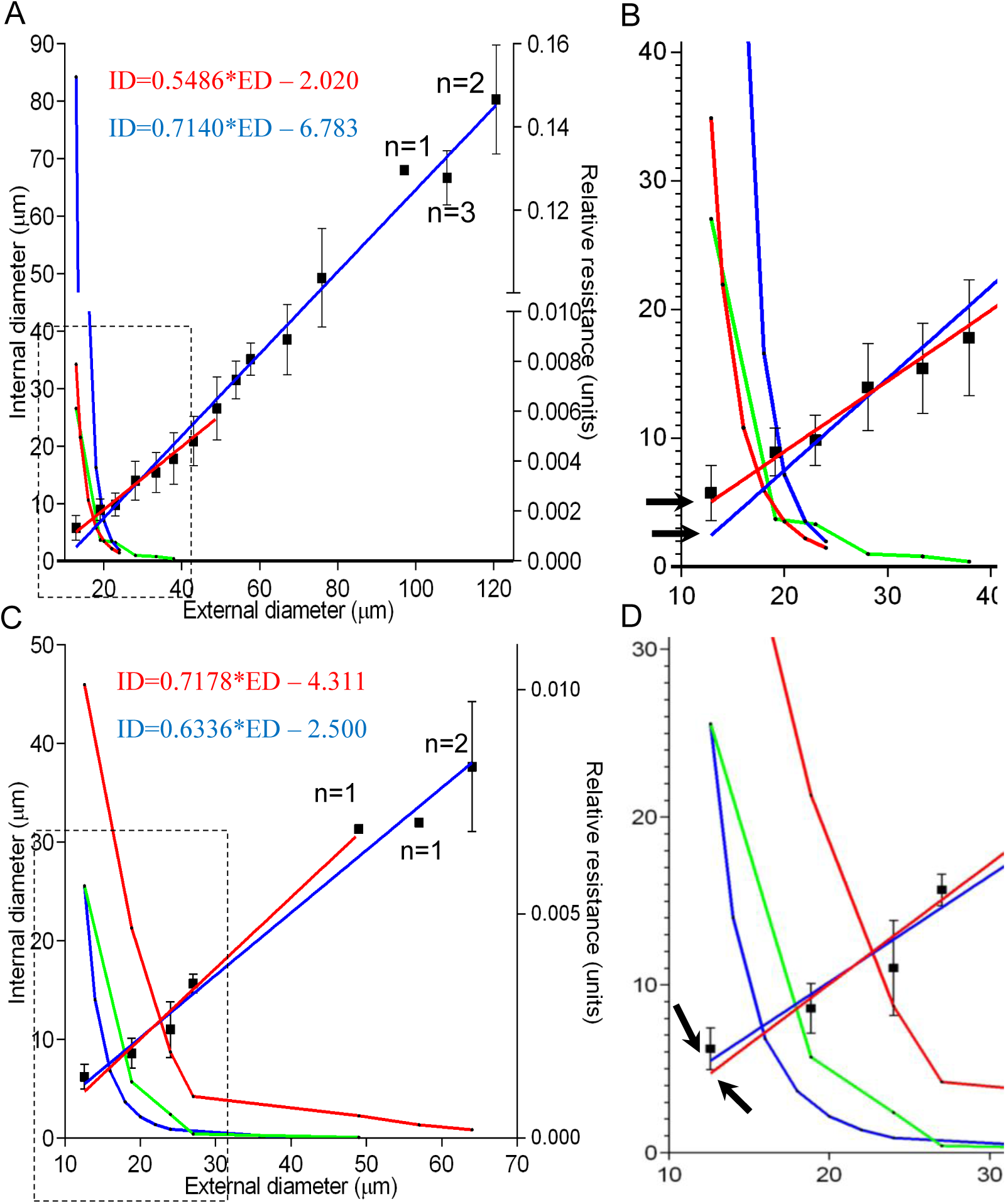
The equations must be verified to avoid the effect of fewer data points for larger vessels. (A) Heart. Means of intervals were approximated with two equations to compute RR curves. The first (red formula and lines) accounted for vessels of ED 10-50 μm. The second (blue formula and lines) accounted for all vessels found (ED=10-120 μm), with only 1-3 values per interval for the largest TAs. The equations were highly distinct between each other (P<0.001), and every line and curve had the highest goodness of fit coefficient *r*^2^ and *R*^2^ ≈0.98-0.99. (B, as boxed area in A) For the smallest arteries, the difference in ID of 1-2 μm (arrows) resulted in a 270% difference in RR curves (P<0.0001). If the RR curve was calculated not from the equation but from means of intervals (green line), it was closer to the red (P>0.88) than to the blue curve (P<0.0001), i.e. the equation for 10-50 μm (red) was correct. (C,D) The same calculation for intestine. The equation for 10-70 μm was correct (P>0.90 vs P<0.22). Points on the lines are mean ± SD for 5-μm ED intervals, n – number of measurements per interval

#### The method enables comparison between species

Using this approach, we compared TAs between normal rat and mouse. While averaged dimensions demonstrate no substantial differences, the complex profiles, linear regression equations and RR curves revealed specific patterns for each species (**Fig 12**). In rats renal TAs have thicker media and narrower lumens compared to mice, but lower resistance, which is consistent with hemodynamic data[74].

**Fig 12.**
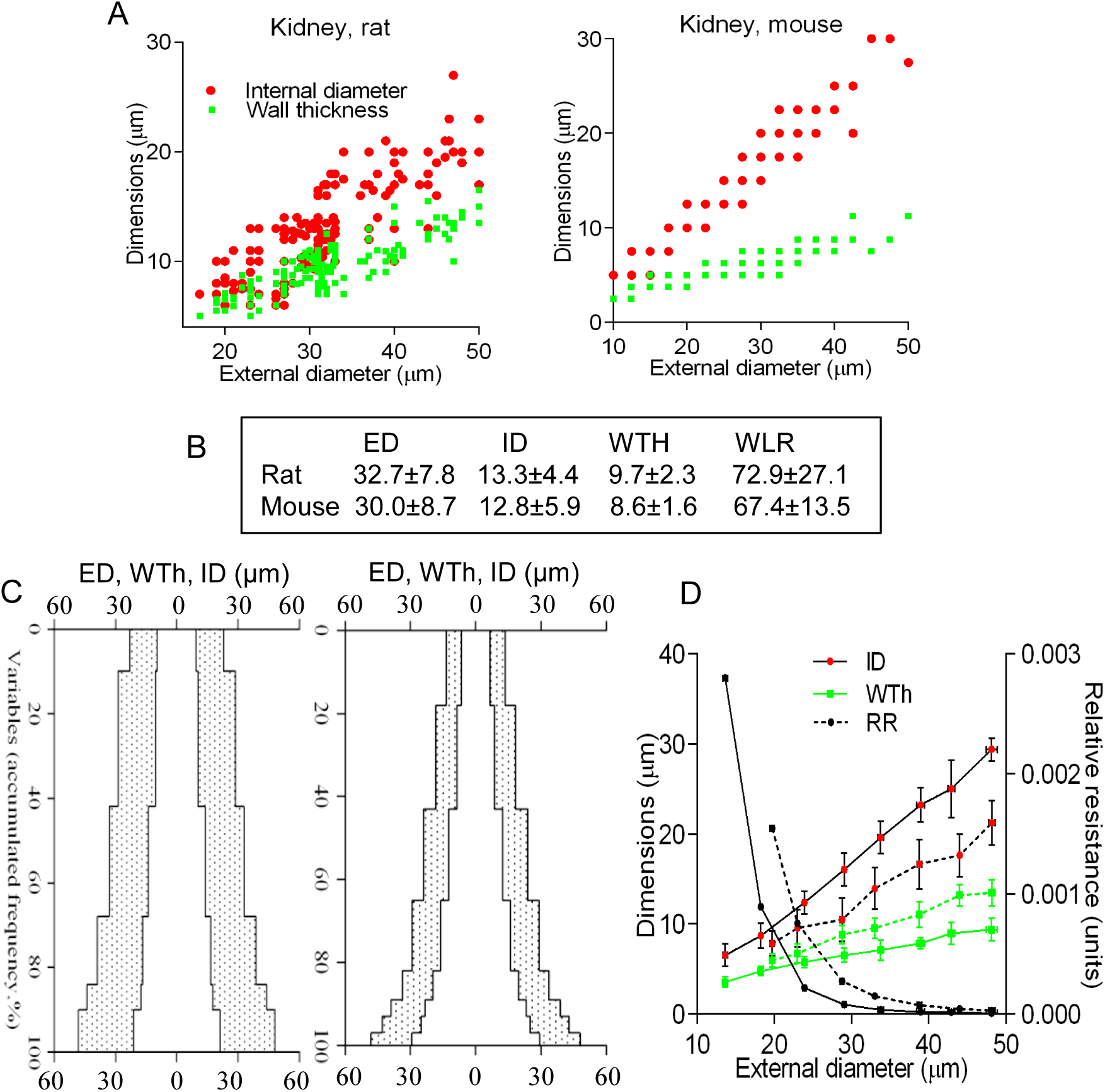
The algorithm enables comparison between species. (A) Scatterplots for ED, ID, and WTh for the rat and mouse. (B) Mean values from scatterplots revealed no difference (P>0.40), including WLR. The corresponding complex profiles (C), and regression equations with the relative resistance curves (D) were very distinctive (P<0.001) for the rat (dashed line) vs mouse (solid line).

### Part II: recognition of remodeling patterns in hypertensive rats on histological sections

#### Conventional measurements exhibit different remodeling patterns in hypertensive rats

At 60 days, the systolic blood pressure in the sham-operated and hypertensive rats was 115±6 mmHg and 217±21 mmHg, respectively (P<0.001). Morphometry data from multiple organs were first analyzed using conventional measures of mean ED, ID and WTh for the arterial ED interval of 10-50 μm. 1K1C hypertension significantly affected TAs in every organ (**Table 2**).

**Table 2.** The spectrum of remodeling variants in terminal arteries of ED 10 – 50 μm.

| **Organ** | **Sham rats** |  |  | **1K1C rats** |  |  | **WLR** | **MCSA** | **NC** |
| --- | --- | --- | --- | --- | --- | --- | --- | --- | --- |
|  | **ED,**<br>**$\mu\text{m}$** | **ID,**<br>**$\mu\text{m}$** | **WTh,**<br>**$\mu\text{m}$** | **ED,**<br>**$\mu\text{m}$** | **ID,**<br>**$\mu\text{m}$** | **WTh,**<br>**$\mu\text{m}$** |  |  |  |
| **Brain** | 23±7 | 13±4 | 5±2 | 19±8**↓ | 7±4**↓ | 6±2 *↑ | ↑ | ↓ | **3-4cI** |
| **Kidney** | 33±8 | 13±4 | 10±2 | 30±9**↓ | 10±4**↓ | 10±3 = | ↑ | ↓ | **3-4b** |
| **Heart** | 27±9 | 13±6 | 7±3 | 27±10 = | 11±4**↓ | 8±3*↑ | ↑ | ↑ | **2-4c** |
| **Pulmo-**<br>**nary** | 29±8 | 21±8 | 4±1 | 33±9**↑ | 22±6 ↔ | 6±1**↑ | ↑ | ↑ | **1-5c** |
| **Skin** | 20±9 | 8±4 | 6±3 | 23±9**↑ | 8±3 = | 7±3**↑ | ↑ | ↑ | **1-5c** |
| **Skeletal**<br>**muscle** | 20±7 | 9±4 | 5±3 | 21±10 = | 8±3**↓ | 6±3 **↑ | ↑ | ↑ | **2-4c** |
| **Bron-**<br>**chial** | 37±9 | 18±7 | 9±1 | 32±10**↓ | 13±6**↓ | 10±2**↑ | ↑ | ↓ | **3-4cI** |
| **Stomach** | 25±7 | 13±5 | 6±3 | 27±11*↑ | 10±5**↓ | 8±3**↑ | ↑ | ↑ | **1-4c** |
| **Intestine** | 19±8 | 9±6 | 5±1 | 24±10**↑ | 9±4 = | 7±3**↑ | ↑ | ↑ | **1-5c** |
| **Adrenal** | 22±8 | 9±3 | 6±2 | 24±11**↑ | 14±5**↑ | 5±3*↓ | ↓ | ↓ | **1-6aII** |
| **Liver** | 22±9 | 8±4 | 6±2 | 22±12 = | 7±4 = | 7±3 = | = | = | **2-5b** |

Data are mean ± SD. \*P<0.05; \*\*P<0.01 NC, numerical classification. ↑ ↓ = indicate increase, decrease or no change respectively vs control values.

To date, the majority of studies use the widely accepted classification[11] (**Fig 13**). However, the obtained remodeling varieties did not fit into conventional definitions. The classification considers ‘inward’ or ‘outward’ as only simultaneous ED-ID decrease or increase (**Fig 13**). The brain, bronchial and kidney should be named ‘inward hypotrophic’ due to reduced ED, ID and MCSA, but that term does not mark increased or stable WTh. The classification has no ‘inward-outward hypertrophic’ type that appeared in means for the stomach. The heart and skeletal muscle should be defined as ‘inward’ because mean IDs were reduced, but constant mean EDs are not recognised, there is no ‘inward-only hypertrophic.’ The same for constant IDs in the pulmonary, skin and intestine, that should be classified as ‘ourward-only hypertrophic.’ For unknown reason, the remodeling had been considered only as keeping constant MCSA, and not constant WTh. The results were based on averaged linear dimensions, and hypertensive values demonstrated a high degree of variability and abnormal distribution (**S4 Fig**). Accordingly, conventional averaging of dimensions led to the designation of diverse remodeling patterns.

**Fig 13.**
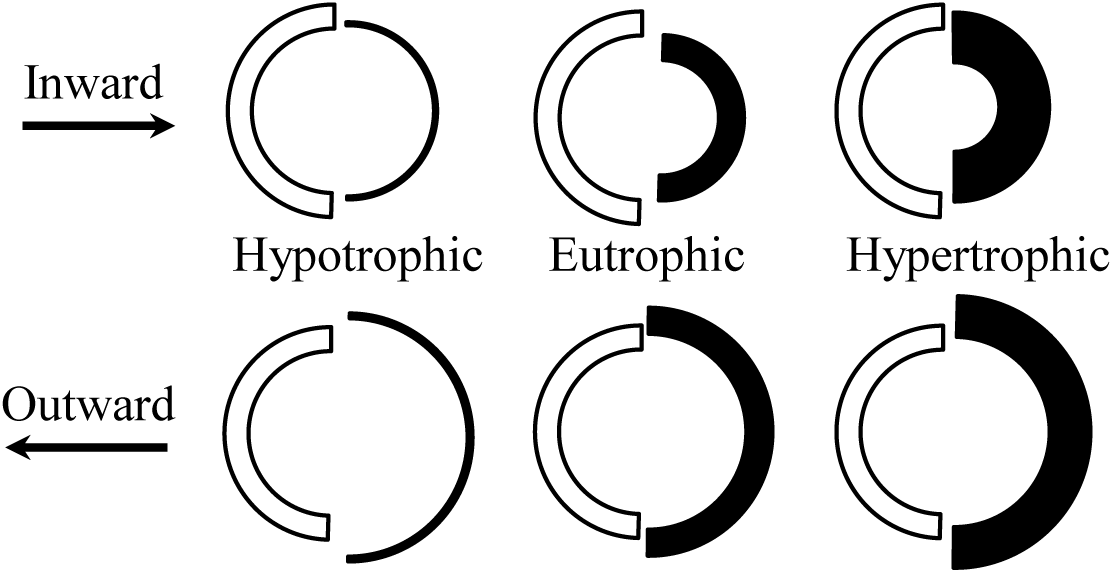
The descriptive conventional classification[11]. The ‘inward’ or ‘outward’ means reduction or increase in lumen; the ‘hypertrophic,’ ‘eutrophic,’and ‘hypotrophic’ indicates increase, no change, or decrease in wall cross-sectional area, respectively.

There is a general assumption that in humans and animals, regardless of the type of hypertension, small arteries develop inward eutrophic or inward hypertrophic remodeling[7],[24]. Other studies have suggested that inward eutrophic remodeling occurs in essential hypertension, while secondary hypertension is associated with hypertrophic remodeling[4]. The concept of uniform remodeling is logical and reasonable since arteries throughout the body have similar structure and regulatory mechanisms. Nevertheless, studies using either random sections or *in vitro* myography present an entire spectrum of possible remodeling patterns. Differences in animal age, heterogeneous post-mortem arterial contraction, variable perfusion pressure, and different histoprocessing and morphometry techniques may account for data inconsistency[3],[47]. However, our attempt to find uniform remodeling in 1K1C rats by using the same tissue preparation and morphometry technique simultaneously in ten organs failed, and instead revealed distinctive, and even opposite patterns (**Table 2**).

Many authors have presented data avoiding classification[14],[75],[76],[77]. Reviews on this subject have focused on the method of tissue preparation and the potential for sampling bias, and the general approach has not been revised[3],[24],[25],[31]. Indeed, the conventional classification was based on sketch-drawings, with assumptions regarding geometric parameters[11] (**Fig 13**). The classification presumed, for unclear reasons, the simultaneous movement of ED and ID as inward or outward. Such movement would occur if the artery remodeling is considered as simple inflation or deflation for an elastic, homogenous, gel-like structured thick-wall cylinder[78],[79]. However, the arterial wall is categorized as a multilayered, helically arranged, fiber reinforced composite, with independent ED vs ID displacement[80]. Wall remodeling is therefore a multilayered interaction involving hypertrophy[81],[82], hyperplasia[21],[83], apoptosis[81], hyalinosis and fibrinoid necrosis[84], [85] of vascular smooth muscle cells, as well as deposition of extracellular matrix[86],[87]. The extent of these changes varies from inner to outer layers[88],[83],[76]. We also have found that quite often statistical data from studies could not be even assigned to a certain type because statistical casualties break geometry rules. For example, mean ED and ID remains constant but WTh increased[60], mean ID decreased while MCSA and WTH remain constant[89].

#### The conventional classification is unable to categorize remodeling variants

In order to precisely categorize remodeling variants in consideration of the conventional designations of inward-outward and hypertrophic-hypotrophic (**Fig 13**), possible conformations of a blood vessel were modeled with the approximation of arteries as thick-wall cylinders capable of changing ED and ID independently (**Fig 14**).

**Fig 14.**
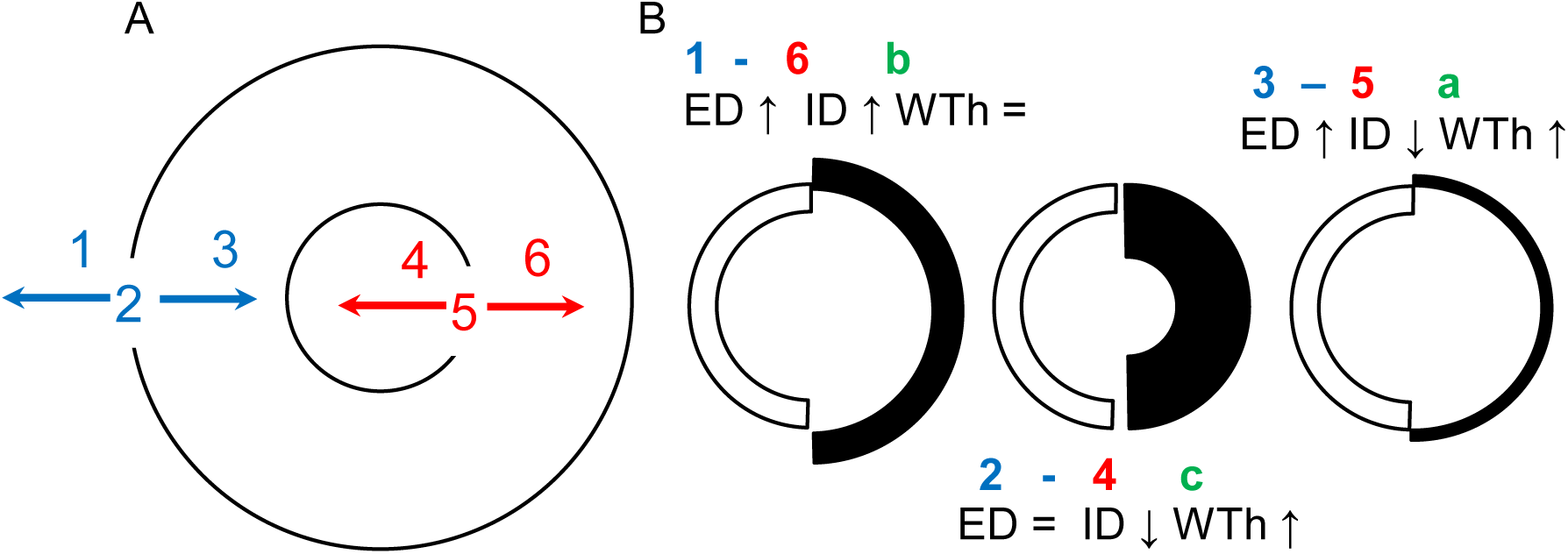
Numerical modeling approach. (A) For numerical modeling the ED (2) and ID (5) changes independently. ED could moves outward (vector 1) to increase ED or inward (vector 3) to reduce ED. ID could shift inward (vector 4) to decrease ID or outward (vector 6) to increase ID. (B) An example of numerical classification. The number 1, 2 or 3 indicates increased (↑), stable (=), or reduced (↓) ED respectively. The number 4, 5 or 6 reflects decreased, constant or increased ID. The letter a, b or c identifies diminished, constant or increased WTh. White semicircles – control, black – predicted remodeling variants.

In a simulation procedure, each variant was considered distinct if a value for only one dimension in the set of five (ED, ID, WTh, MCSA, WLR) was different from the value for another set. The numerical modeling (**Fig 15**) revealed not six, as empirically suggested, but twenty variants of arterial remodeling. For example, increasing ED produces eight (1-4, 1-5, 1-6aI, 1-6aII, 1-6b, 1-6 cI-cIII) possible variants by combination of changing ID, WTh and MCSA in different directions. A constant ED results in two variants (2-4, 2-6), and decrease in ED generates nine variants (3-4 aI-aIII, 3-4b, 3-4 cI-cIII, 3-5, 3-6). Variants 1-6 cI-III, 3-4 aI-III and 3-4 cI-III are possible because not only the direction but even gradients of change between ED and ID could elicit unique remodeling patterns, which were impossible to classify with conventional definitions.

**Fig 15.**
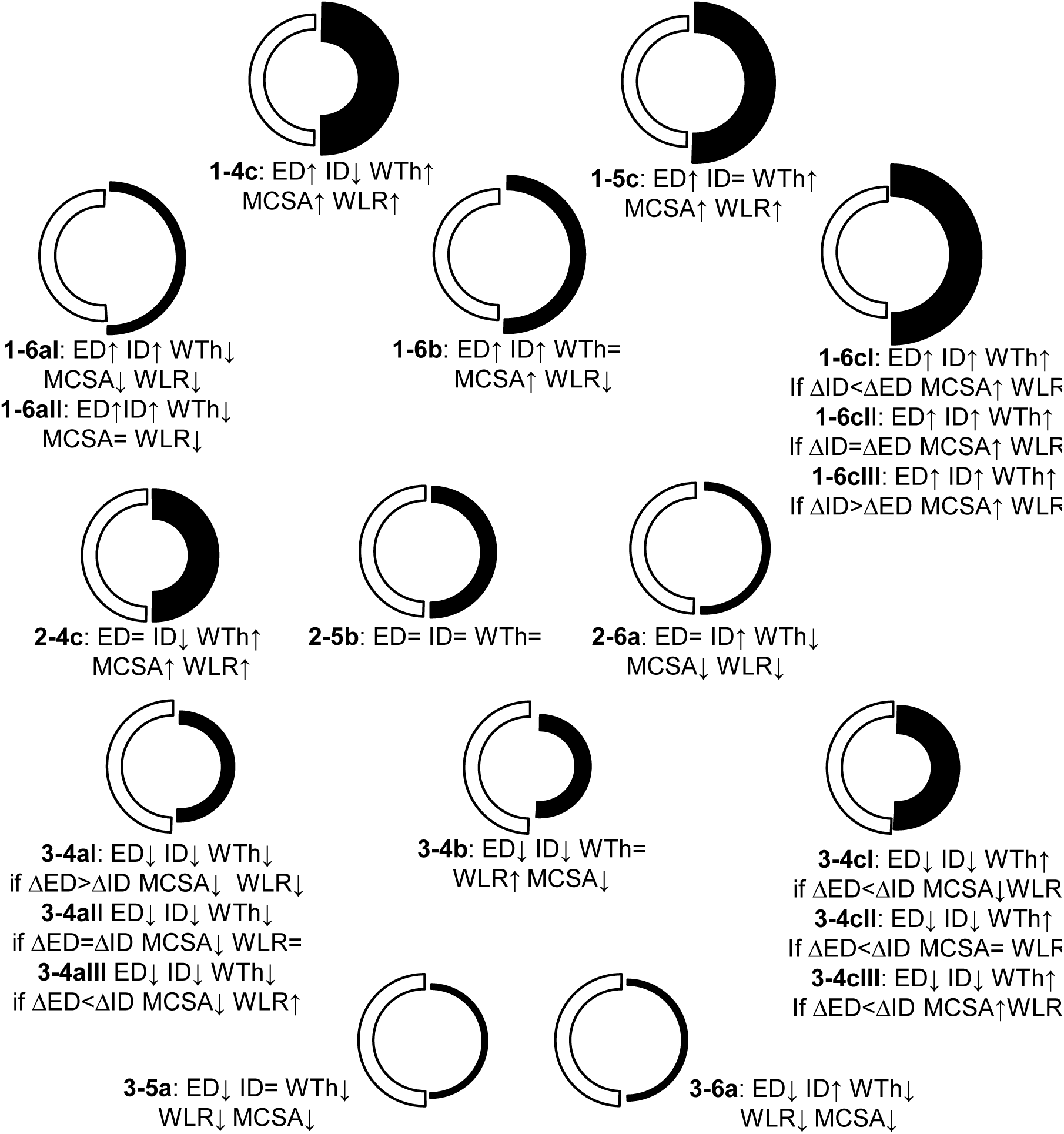
The formalization of numerical modeling for predicted remodeling variants. The number **1**, **2** or **3** indicates increased (↑), stable (=), or reduced (↓) ED. The number **4**, **5** or **6** corresponds to decreased, constant or increased ID. The letter **a**, **b** or **c** identifies increased, constant or diminished WTh respectively, for the same set of ED and ID. The additional Roman numerals **I**-**II**-**III** detail the dynamics in MCSA and WLR for the same set of ED, ID, and WTh due to possible gradients in ED vs ID displacement. The 2-5b variant indicates no remodeling. White semicircles – control, black – predicted remodeling variants.

Therefore, the conventional term “hypertrophic” would equally apply to nine variants: 1-4, 1-5, 1-6cI-III, 2-4, and 3-4cI-III. The term “outward” would cover simultaneously 1-4, 1-5, and all six sub-variants 1-6. The term “eutrophic” would equally apply to 1-6b, 3-4b or 3-4cII, while “hypotrophic” could be true for 1-6aI-II, 2-6, 3-4aI-III, 3-5, and 3-6. According to this numerical classification, in 1K1C rats TAs demonstrated seven remodeling variants across organs (**Table 2**). Studies using either random sections (**Table 3**) or *in vitro* myography (**S3 Table**) are also very inconsistent.

**Table 3.** Numerical classification applied to arteries studied on random histological sections.

| References | Experimental model | Arterial location | NC |
| --- | --- | --- | --- |
| Liu et al.[17] | 2K1C (PF) | renal arterioles 30-60µm<br>> 60µm | 2-4c<br>2-5b |
| Helmchen et al.[44] | 2K1C (IF) | interlobular | 1-5c |
| Qin et al.[82] | 2K1C(PF) | mesentery | 1-4c |
| Korsgaard et al.[90] | 1K1C, unfixed,<br>wire myograph | mesenteric III branch<br>renal arcuate arteries | 3-4cIII<br>2-5b |
| Deng et al.[77] | wire myograph, 2K1C<br>1K1C | mesentery III order<br>mesentery III order | 3-4cIII<br>3-4cII |
| Zhou et al.[87] | 2K1C, wire myograph | renal arteries | 3-4cIII |
| Kinuno et al.[13] | SHR (PF)<br>SHR+uninephrectomy (PF) | renal interlobular<br>renal interlobular | 1-5c<br>1-6aII |
| Owens et al.[18] | SHR (PF) | mesentery, I branch<br>II branch<br>III - IV branch | 3-4cII<br>2-5b<br>3-4cIII |
| Smeda et al.[16] | SHR (PF) | renal, ID~20-60µm<br>renal, ID~300-60µm | 2-5b<br>1-5c |
| Johansson B.[91] | SHR (PF)<br>SHRSP (PF) | cerebral extracranial<br>cerebral intracranial<br>cerebral extracranial<br>cerebral intracranial | 1-6cI<br>1-5c<br>1-6cIII<br>1-6aII |
| Nordborg et al.[12] | SHR (IF)<br>SHRSP (IF)<br>SHR (IF)<br>SHRSP (IF) | IF, mesentery<br>mesentery<br>renal<br>renal | 3-4cIII<br>3-4cII<br>1-5c<br>1-6cI |
| Leh et al.[14] | SHR (IF) | renal afferent arterioles | 1-6cI |
| Ohara et al.[15] | SHR (PF) | PF, renal interlobular | 2-4c |
| Kost et al.[92] | SHR (PF) | renal afferent<br>interlobular<br>arcuate | 2-5b<br>2-5b<br>3-5a |
|  | SHR+AngII (PF) | renal afferent<br>interlobular<br>arcuate | 2-5b<br>1-5c<br>1-5c |
| Lee et al.[47] | SHR (PF) | mesentery,<br>ID~120-250 µm | 1-5c |
| Limas et al.[50] | SHR, age dependent (PF)<br>renal, ED~50-100 µm | 10 weeks | 2-5b |
|  |  | 20 weeks | 1-5c |
|  |  | 28 weeks | 3-4b |
|  |  | 48 weeks | 2-4c |
| Casare et al.[19] | AngII (PF) | renal afferent<br>interlobular | 1-6cII<br>1-6aII |
| Berry et al.[21] | DOCA (PF)<br>intrarenal, | ED ~ 400-500µm<br>ED~ 200-300µm | 1-6cI<br>1-5c |
| Mazzali et al.[40] | Hyperuricemia (IF) | renal afferent arterioles | 1-4c |

2K1C, two kidney-one clip; Ang, angiotensin; DOCA, deoxycorticosterone acetate; IF, immersion fixed; PF-perfusion fixed; SHRSP – spontaneously hypertensive rats stroke prone; NC – numerical classification.

We have found only one study that applied a numerical approach to remodeling[31]. The numerical classification proposed here unambiguously defines any wall conformation, and could be applicable to remodeling not only in hypertension but diabetes, atherosclerosis[93],[94], high or low blood flow[48],[27], restenosis[95], vasculitis[96], or bronchial remodeling[97].

#### Remodeling patterns are not identified with WLR, RI or GI

To decide whether remodeling varieties (**Table 2, 3 and S3 Table**) truly exist or are derived from conventional averaging, we tested methods recognizing that arteries follow one of twenty predicted variants.

Increased or decreased WLR is widely used as the main criteria of ‘inward’ vs ‘outward’ wall conformation[3],[4],[5]. It is not appropriate because WLR could be similarly increased in nine, decreased in eight, and unchanged in three variants. Therefore the frequently used WLR could unambiguously define neither wall thickening nor lumen narrowing (**Fig 15**).

RI and growth index GI are also regarded as the main parameters to estimate remodeling[35],[98],[24]. According to the primary data[6], basilar arteries in SHR developed the variant 3-4cIII as a combination of hypertrophy and rearrangement (remodeling) of vascular smooth muscle cells (**Fig 16**). The proposed combination was explained with two formulae. The formula (1) below was based on the first presumption: what would the ID be if vessels underwent the variant 3-4cII (ID*3-4cII*) i.e. hypertensive MCSA (MCSA*hr*) remains equal to normal MCSA (MCSA*n*)? The formula (2) below was based on the second presumption: what would the ID be if vessels developed the variant 2-4c (ID*2-4c*) i.e. hypertensive ED (ED*hr*) remains equal to normal ED (ED*n*)? To create the formula (1), the larger MCSA*hr* was ignored in favor of a proposed MCSA*n*=MCSA*hr* to calculate percent of encroached lumen if the variant 3-4cII would occur:

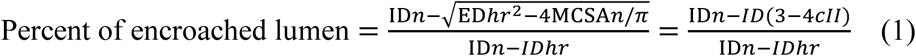

**Fig 16.**
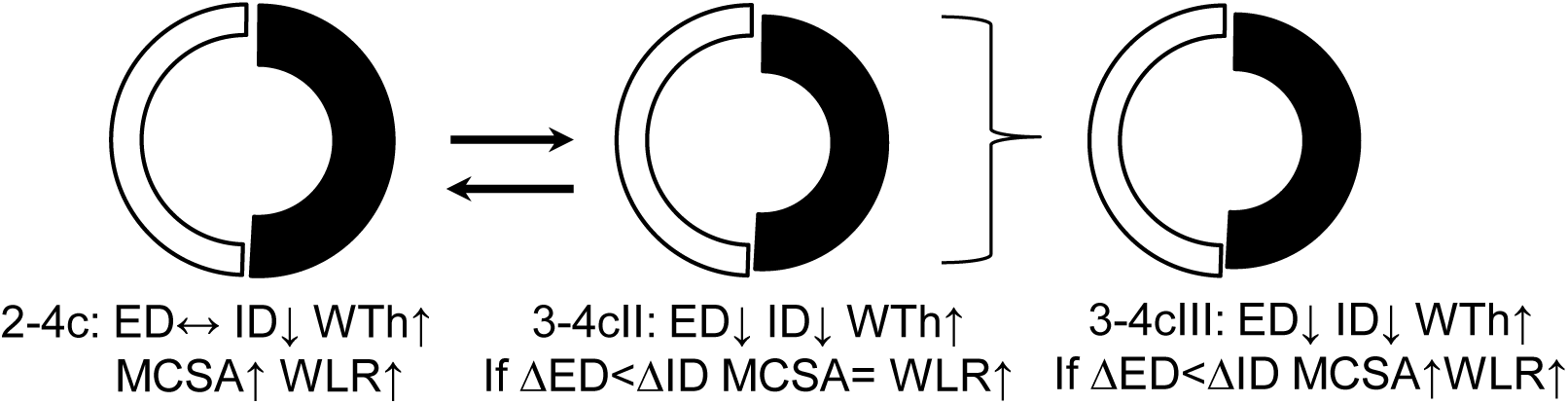
The first attempt to quantify remodeling[6]. The reductions in both mean ED and ID were interpreted by the authors as a possible combination of hypertrophy (variant 2-4c) and rearrangement of vascular smooth muscle cells (variant 3-4cII). White semicircles – control, black – predicted remodeling variants

For the formula (2), the smaller ED*hr* was ignored in favor of a proposed ED*n*=ED*hr* to calculate percent of encroached lumen if the variant 2-4c would occur:

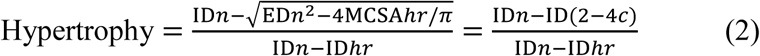

The primary definition “percent of encroachment lumen” in the first formula was renamed the “remodeling index”, and the second formula for “hypertrophy” was modified to “growth index” by[98]. Unlike the second formula, GI counted true MCSAs:

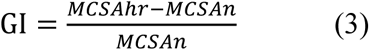

Accordingly, the formulae were intended to quantify only *<u>a subjective interpretation</u>* of 3-4cIII as a combination of 2-4c and 3-4cII but that interpretation could be valid as any combination between eight variants with diminished ID (2-4c, 3-4aI-III, 3-4b, 3-4cI-III). Therefore calculation of RI is also incorrect because the formulas were designated to analyze only the proposed combination (**Fig 16**).

GI does not bear any information about growth, as it simply counts the percentage change in MCSA that could be assigned to different variants (**Fig 15**). This may account for the variable RI and GI values found in *in vitro* myography experiments, which are considered gold standard due to sampling of similar mesenteric arteries branches. According to formulae (1) and (2), the sum RI+GI must be 100%, but it appears only in the first authors’ publication (**S3 Table**). The question still remains: how to recognize a particular remodeling variant?

#### Remodeling patterns are not identified on 2D graphs

The analysis of the simple annular ring in histological sections represents a conundrum, since arteries have no ‘reference points’ from which they have been modified to assume a remodeled shape (**Fig 17**). According to the literature and own data, any of the 20 predicted types of remodeling could occur (**Table 2, 3 and S3 Table**).

**Fig 17.**
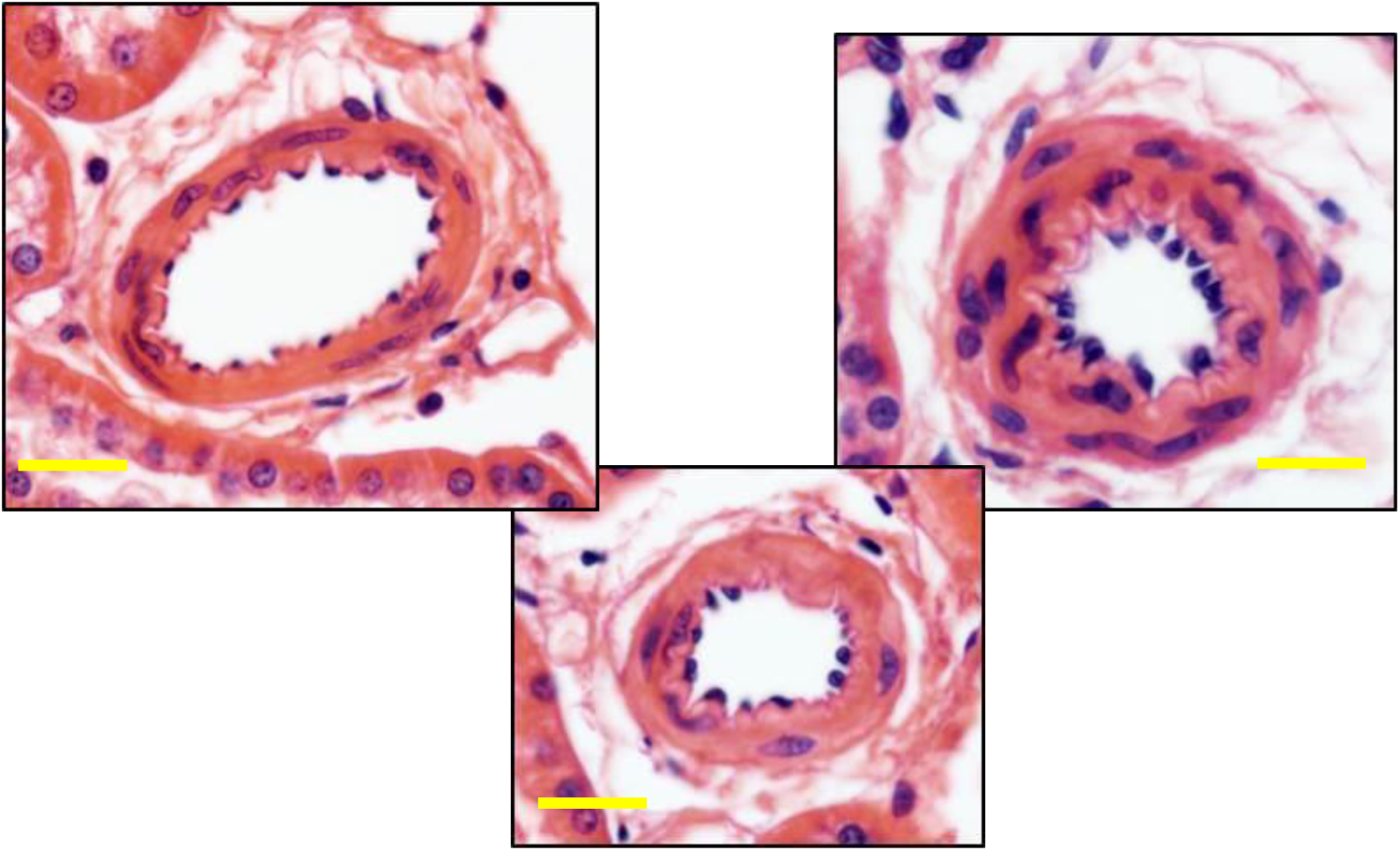
The conundrum in arterial remodeling assessment. Each artery could be an image of another one before or after remodeling: there is no “reference point” indicating its previous dimensions. Scale 10 μm.

As in control rats, primary measurements were organized in 5 μm intervals, the mean value for each interval was calculated, and the complex profile and linear regression equations built from the means of intervals (**S4 Table**). Data analysis revealed that comparison of complex profiles also could not be used to identify remodeling variants because it was complicated by the existence of uncertain starting points for superimposition of a hypertensive graph on the control one (**Fig 18A, S5 Fig**). Furthermore, there are no data indicating how vessel remodeling progresses in a length-wise fashion, except for the presence of increased vessel tortuosity[29],[63]. While the common starting points were unclear, varying accumulated frequencies suggested distinct remodeling patterns even in different segments within the same organ (**Fig 18A, S5 Fig**).

**Fig 18.**
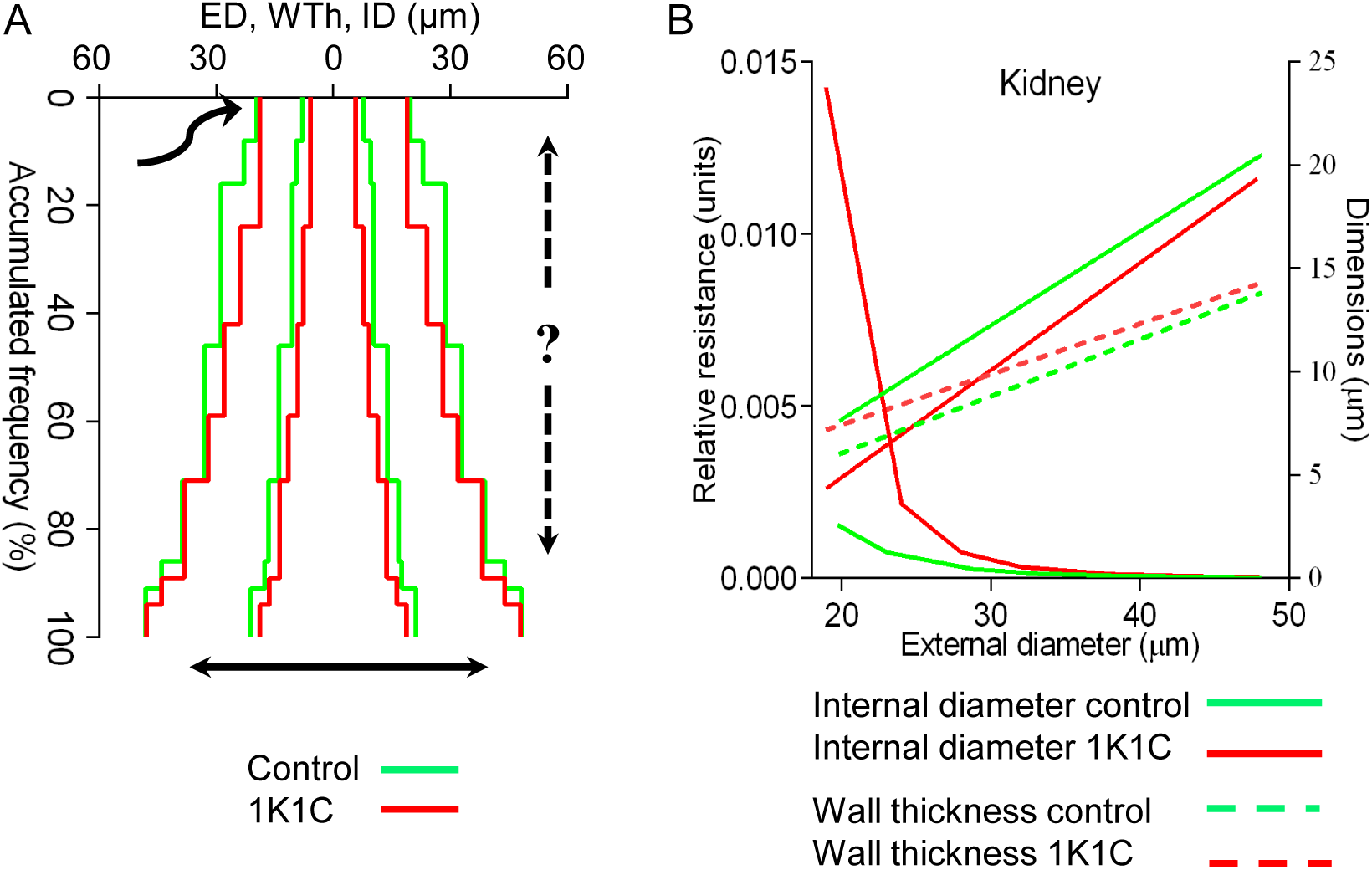
Considerations for arterial remodeling assessment. (A) The profile of hypertensive (red) renal TAs was superimposed on the control (green). The starting point (the curved arrow) is uncertain: inward–outward remodeling (the solid double arrow) is well known, while proximal-distal shifting (the dashed arrow) is unacknowledged. Uneven accumulated frequencies displaced interval values irregularly. Complementary graphs are in **S5 Fig.** (B) Linear regression lines of renal TAs demonstrated decreased ID (solid lines) and increased WTh (dashed lines) in hypertension. The slopes were different (P<0.0001). Complementary graphs are in **S6 Fig.**

Next, we tested if linear regression could identify remodeling patterns. Compared to the complex profiles, the linear regressions demonstrated smaller IDs and bigger WThs for the same ED similarly in all organs, except the kidney and adrenal (**Figure 18B, S6 Fig**).

To understand how shifts in regression lines could verify remodeling, every predicted variant, for each organ, was simulated from control linear regression equations (**S4 Table**). Some studies considered that not WTh/ID but other ratios such as EP/IP[42],[41], or MCSA/LCSA[13],[38],[45], or MCSA/IP[22] are more reliable and informative. Therefore we tested all possible dimensions and their ratios to demonstrate the lack of reliability in 2D graphs. Simulation was done for: linear sizes including ED, ID, WTh, internal perimeter (IP), external perimeter (EP); areas, such as MCSA, lumen cross sectional area (LCSA), total area (MCSA+LCSA), and ratios, including WLR, EP/IP, and MCSA/LCSA.

For all tested dimensions, the regression line displacement from the control line was similar for as many as 11 variants (**Fig 19, S7 Fig**). Any linear size, area or their ratio exhibited similar displacement for many different remodeling variants. While the use of linear regression equations can verify the presence of remodeling, the equations are unable to define a specific remodeling pattern. Thus, it was not possible to distinguish arterial spatial conformations on routine 2D graphs.

**Fig 19.**
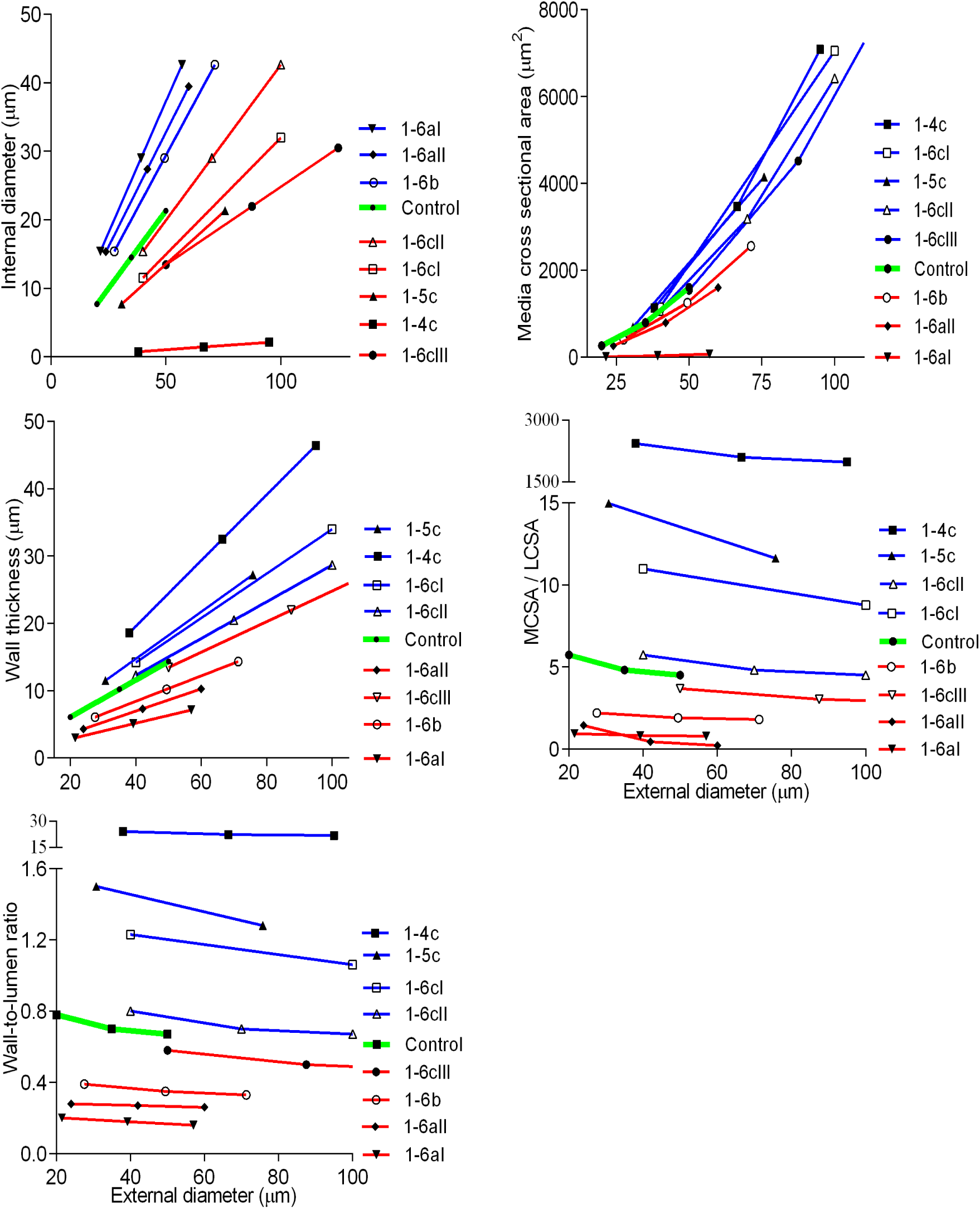
Remodeling variants are not distinguishable in two-dimensional graphs. The simulated remodeling of renal arteries for the variants 1-4c, 1-5c and all 1-6. For any parameter a line shift was similar to others, and could distinguish only increase or decrease from control but not verify a particular pattern. MCSA - media cross sectional area, LCSA – lumen cross sectional area. Complementary graphs are in **S7 Fig.**

#### A method of 3D-modeling simulation recognizes remodeling type

A 3D-modeling simulation was applied to address the limitations with 2D graphs. There were four steps to this method. For example, for the brain, IDs and WThs were calculated from the control linear regression equations (**S4 Table**) and placed in 3D-graphs (**Fig 20, S8 Fig**). Then, IDs and WThs for each of twenty possible remodeling variants were computed from the control equations and added to the 3D graphs. Every remodeling pattern was simulated by calculating its possible maximal deviation in dimensions that were arbitrarily limited to 300% for wall thickening or lumen widening, and 99% for wall thinning or lumen narrowing. IDs and WThs were then calculated from the hypertensive regression equations (**S4 Table**) and also added to 3D graphs. Finally, line proximity between real and predicted values was verified with methods of analytic geometry in 3D space[99]. From all possible variants the hypertensive remodeling in cerebral TAs corresponded only to the variant 2-4c.

**Fig 20.**
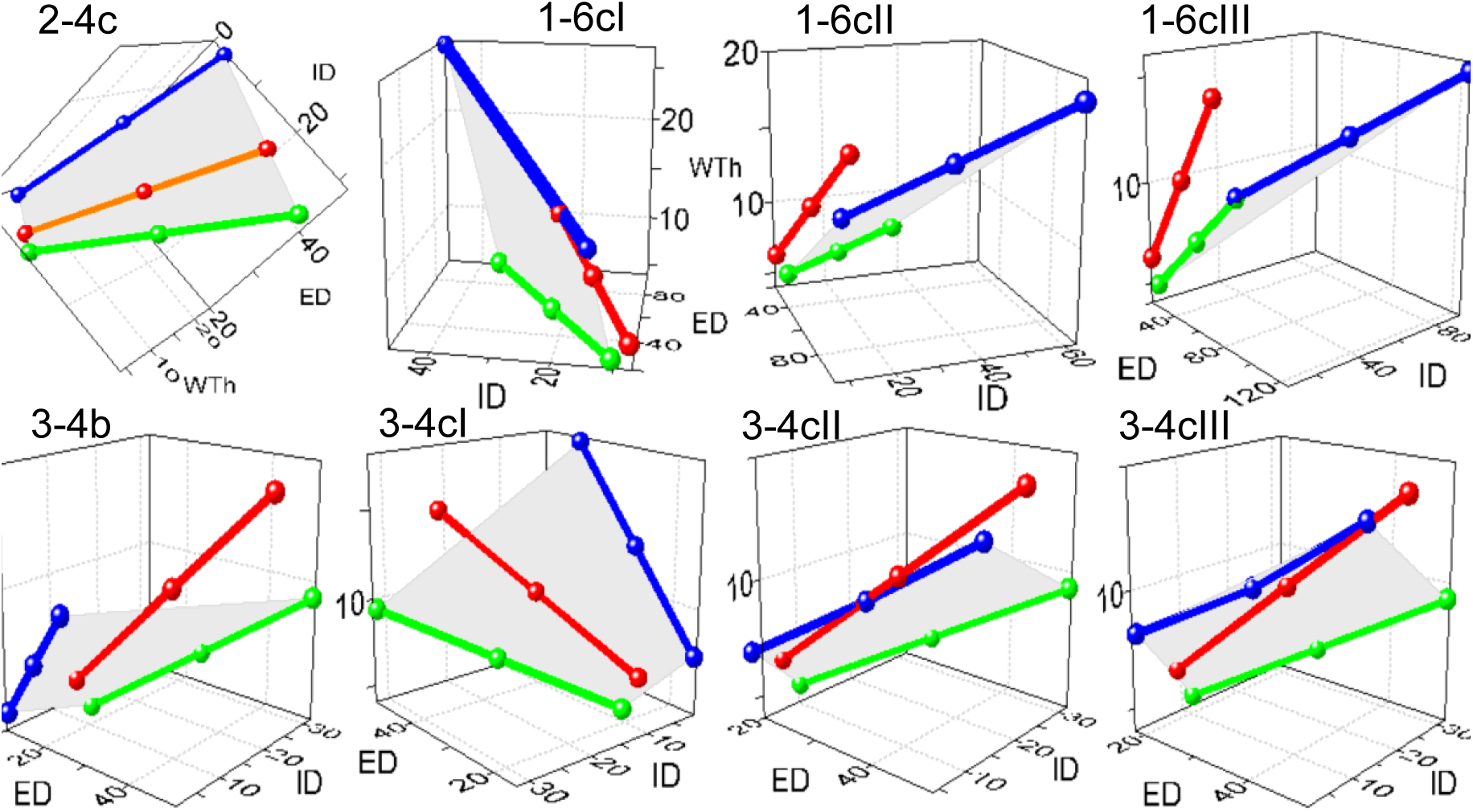
3D modeling simulation for the brain. In 3D graphs, the control equation (green lines) and each predicted variant (blue lines) are connected by the planes (grey) representing sets of all possible values that ED, ID, and WTh could acquire during transformation. The red lines represent the regression equation for hypertensive rats (**S4 Table**). The red lines were not congruent with any of the predicted planes, except for the variant 2-4c. The method readily distinguishes the gradients ΔED, ΔID, and ΔWTh for the variants 1-6 and 3-4 that could only be implied in 2D graphs in Fig 19. Axis X – ED, axis Y – ID, axis Z – WTh, μm. Complementary graphs are shown in **S8 Fig., S19 and S20 Video files.** Throughout the article, coordinates in some graphs were rotated at different angles to optimize display.

The brain, heart, lung, bronchi, liver, stomach, intestine, skin, and skeletal muscle developed the same remodeling variant 2-4c, supporting the concept of uniform remodeling (**Fig 21**).

**Fig 21.**
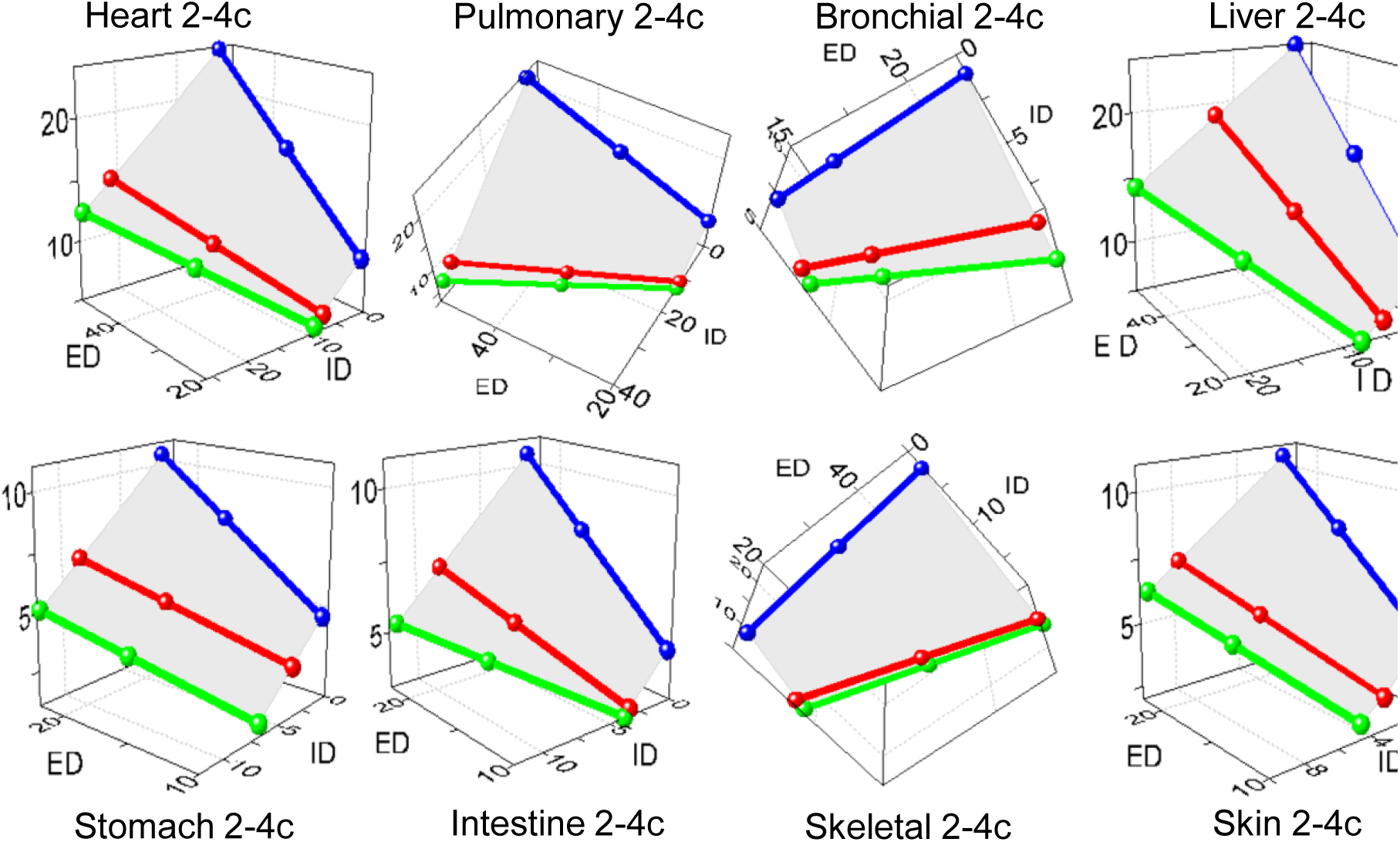
The common remodeling variant in hypertensive rats. The same variant 2-4c was found in most organs.

We also found two novel variants of remodeling, as predicted. The adrenal TAs exhibited more complex spatial rearrangements. The distal segments with ED ~10-20 μm followed the variant 2- 4c, as other organs (**Fig 22A and S9 Fig**). The proximal segments of ED ≈ 30-50 μm transformed oppositely - through the variant 1-6aI with a distended lumen and reduced WTh and MCSA (**Fig 22B and S10 Fig**).. This was evident even on 2D graphs (**S6 Fig**). The adrenals in the 1K1C model experienced a high blood flow. The blood, shunting from clipped or ligated main renal arteries, induced the variant 1-6aI in the proximal segments, i.e. wall distention and thinning, as described for remodeling in high flow models[26],[27],[30]. However, distal TAs demonstrated the same variant 2-4c as in other organs. It may be that smallest arteries (arterioles) are structurally more resistant to distention from high blood flow, due to the absence of elastic laminae and adventitia[100]. Such segmental arterial remodeling has not been verified previously to our knowledge, and indeed was present in TAs with EDs ranging from 10-50 μm.

**Fig 22.**
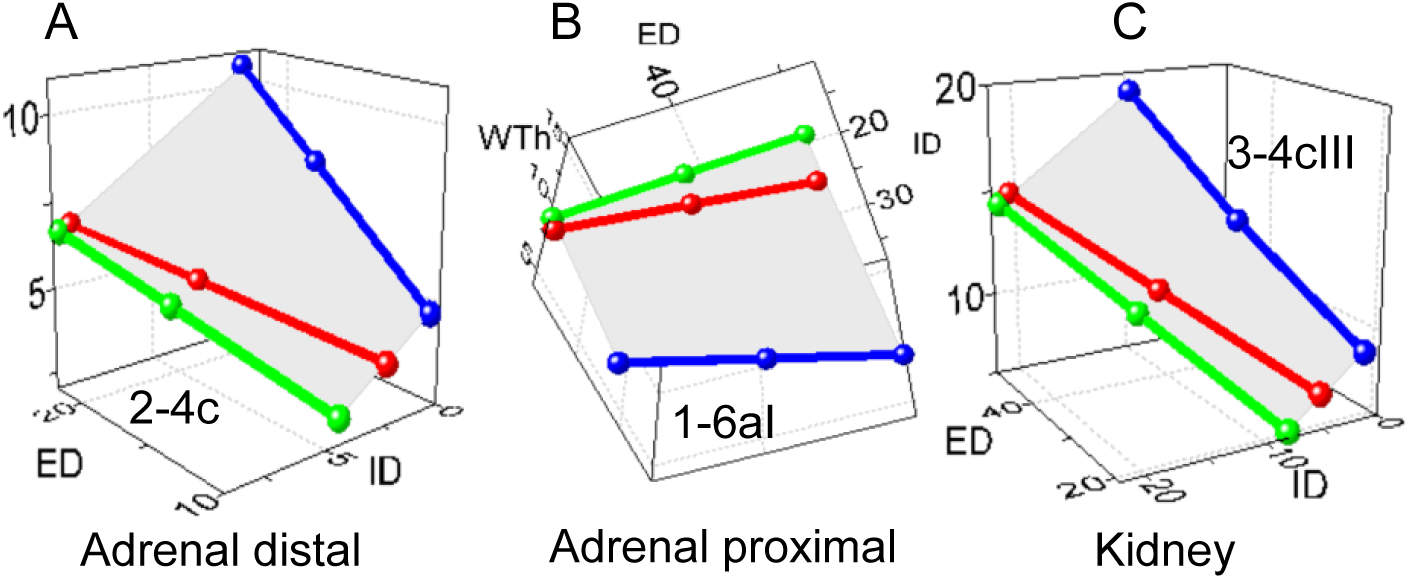
The distinct remodeling variants in hypertensive rats. (A) Adrenal distal segments of ED ~10-20 μm were congruent with the variant 2-4c, as other organs. Complementary graphs are in **S9 Fig.** (B) Adrenal proximal segments of ED ~30-50 μm were congruent with the variant 1- 6aI. Complementary graphs are in **S10 Fig.** (C) The variant 3-4cIII appeared in the kidney. Complementary graphs are in **S11 Fig.**

Renal TAs developed the variant 3-4cIII (**Fig 22C and S11 Fig**), that was responsible for the largest TRR increase. Reduced blood flow in stenotic kidneys[28],[29] should initiate one of the variants (3-4a-b), which has been described for low-flow models[26],[30]. The renal arteries also experienced enhanced RAS activity that likely initiated the variant 2-4c in most organs. Development of the variant 3-4cIII might therefore reflect an interaction between flow-induced and pressure-induced stimuli. We speculated that remodeling by the variant 3-4cIII is a phenomenon of great importance that could be extended beyond the 1K1C model. It is well known that hypertension is combined with atherosclerosis in 70-80% of patients[101]. Therefore TAs in the brain and heart could be exposed to the low flow due to stenotic atherosclerotic plaques in proximal segments of cerebral and coronary arteries, and activated RAS as well. Presumably, TAs would develop the same variant 3-4cIII with the most increased resistance and reduced flow, which might be responsible for lesions in the heart and brain, being recognized as critical target organs.

We have provided the first quantitative evidence that pulmonary arteries respond to the hypertensive stimulus in 1K1C rats, and in the same remodeling pattern as other organs in the systemic circulation. Our data also represent the first evidence that substantial TAs remodeling occurs in the liver, intestine, and bronchi, which are not considered typical targets in hypertension[5],[25].

#### The algorithm enables comparison among different studies

Contradictions of two types have been found in conventional remodeling analysis. First, results do not fit the geometric formulae for annular ring dimensions. We applied the term “statistical artifact” (SA) to indicate them. Second, results could be geometrically correct but showed different, opposite, or no remodeling patterns for the same model, organ, or arterial segments (**Table 2, 3 and S3 Table**). In order to demonstrate that our method could resolve this problem, we used 3D modeling to reanalyze data from studies where linear regression had been calculated, although only few articles could be found. Comparative analysis would not be possible with primary averaged data only.

Two studies claimed unusual non-pressure-related wall thickening after comparing arterial remodeling between aortic coarctation and Goldblatt-induced hypertension in rats[17],[102]. One study[102] demonstrated many SAs (**S5 Table**). After coarctation, means of WThs and WLRs were elevated despite stable EDs, IDs, and MCSAs. In 1K1C rats, stable IDs with greater EDs, WThs, and WLRs pointed to the variant 1-5c, although stable MCSAs did not support this designation. A separate study[17] was also inconsistent (**S6 Table**). In both models, averaged dimensions indicated no remodeling in larger segments, but the variant 2-4c in smaller arteries. In coarctation, SA occurred in arteries of ED interval 31-60 μm: unchanged MCSAs did not correspond to stable EDs with lumen narrowing. In two kidney one clip (2K1C) rats, constant EDs and IDs did not follow MCSA and WTh expansion.

Using primary data[17],[102], we calculated linear regressions and applied 3D-modeling. Aortic coarctation exhibited no remodeling in renal or cremaster arteries (**Fig 23A, S12 and S13 Fig**), i.e. there is no non-pressure related TAs thickening distally to aortic coarctation, in normotensive renal and cremaster arteries, as mean values indicated[102],[17]. No further data have been published to support such normotensive thickening[5],[24]. In contrast, the variant 1- 6aI has been recognized in the liver[48]. Goldblatt hypertension induced the same variant 2-4c in all segments of cremaster and renal arteries (**Fig 23B and 23C, S14 and S15 Fig**). We also used primary data to recalculate morphometry in renal arteries in SHR[39], and found the variant 2-4c (**Fig 23D and S16 Fig**).Accordingly, our data suggest that the variant 2-4c is predominant in hypertensive remodeling. The variant 1-5c (stable lumen with wall thickening) was identified in a clinical case of pulmonary hypertension due to congenital mitral stenosis[55] (**Fig 23E and S17 Fig**). That example underlines the precision of our proposed algorithm, since primary data included only fourteen measurements from one pulmonary biopsy. This type of pulmonary hypertension is developed during organogenesis and intensive arterial vascular smooth muscle cells proliferation[103], which could be associated with a stable lumen and wall thickening.

**Fig 23.**
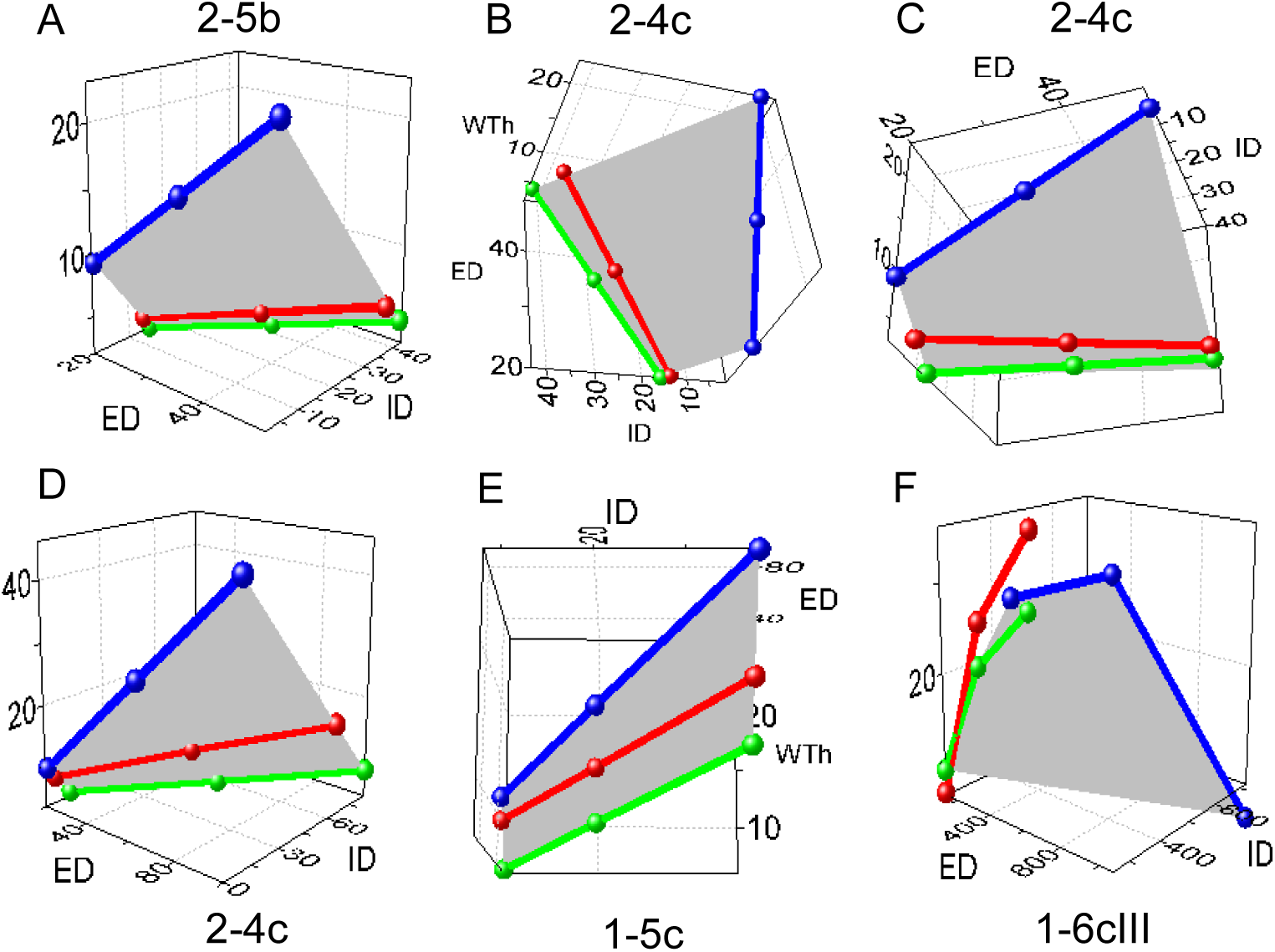
The method enables direct comparison between different studies. (A) Aortic coarctation had no effect on remodeling in renal and cremaster arteries. Complementary graphs are in **S12 and S13 Fig.** (B, C, D) The same variant 2-4c characterized remodeling in cremaster arteries of 1K1C, untouched kidneys of 2K1C rats, and kidneys in SHR respectively. Complementary graphs are in **S14-S16 Fig.** (E) The distinctive variant 1-5c was detected in pulmonary arteries due to congenital mitral valve stenosis Complementary graphs are in **S17.** (F) Data of the subcutaneous arteries was approximated inappropriately with linear regression. Complementary graphs are in **S18 Fig.**

We also tested the linear regression on subcutaneous arteries, reported in patients with essential hypertension[104]. 3D-modeling detected multiple curved lines (**Fig 23F and S18 Fig**) that were the result of an incorrect approximation of the area / length scatterplot by the linear regression. Plotting areas against lengths follows the exponential function[21],[22] (**Fig 19**). The same error we found in other studies[18].

#### Connecting measurements to hemodynamic values

In addition to proposed TRR and TC, it was also desirable to estimate how media volume had changed in remodeling. The terminal media volume (TMV) could be calculated from complex profiles, similar to TC. However, in the current study we found that variation in accumulated frequencies would significantly affect TC and TMV values. The same negative issue for complex profiles to distinguish remodeling patterns was described above (**Fig 18A**). Accordingly, the linear regression equations, established for each organ, with counting ID and ED for the largest (50 μm) and smallest (10 μm) vessel calibers, provided more reliable estimates. TC and TMV were then computed by the formula for the truncated cone volume:

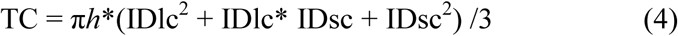

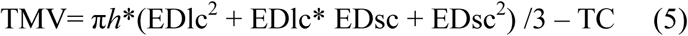

where EDlc, IDlc and EDsc, IDsc were the largest and smallest caliber limits, respectively, and *h* was the range of calibers (50 μm – 10 μm = 40 μm).

Using this calculation, 1K1C hypertension induced a significant 2 - 5-fold increase in TRR in all organs (**Fig 24A**), correlating significantly with data obtained by physiological measures: in the Goldblatt model increased RVR has been shown in every organ, and is most pronounced in kidneys[105],[106],[107],[108]. It is noteworthy that the kidney has the highest increase in TRR (9-fold), while the pulmonary arteries – the lowest. The increased TRR and TC observed in the adrenal glands were due to different remodeling patterns in the proximal and distal arterial segments.

**Fig 24.**
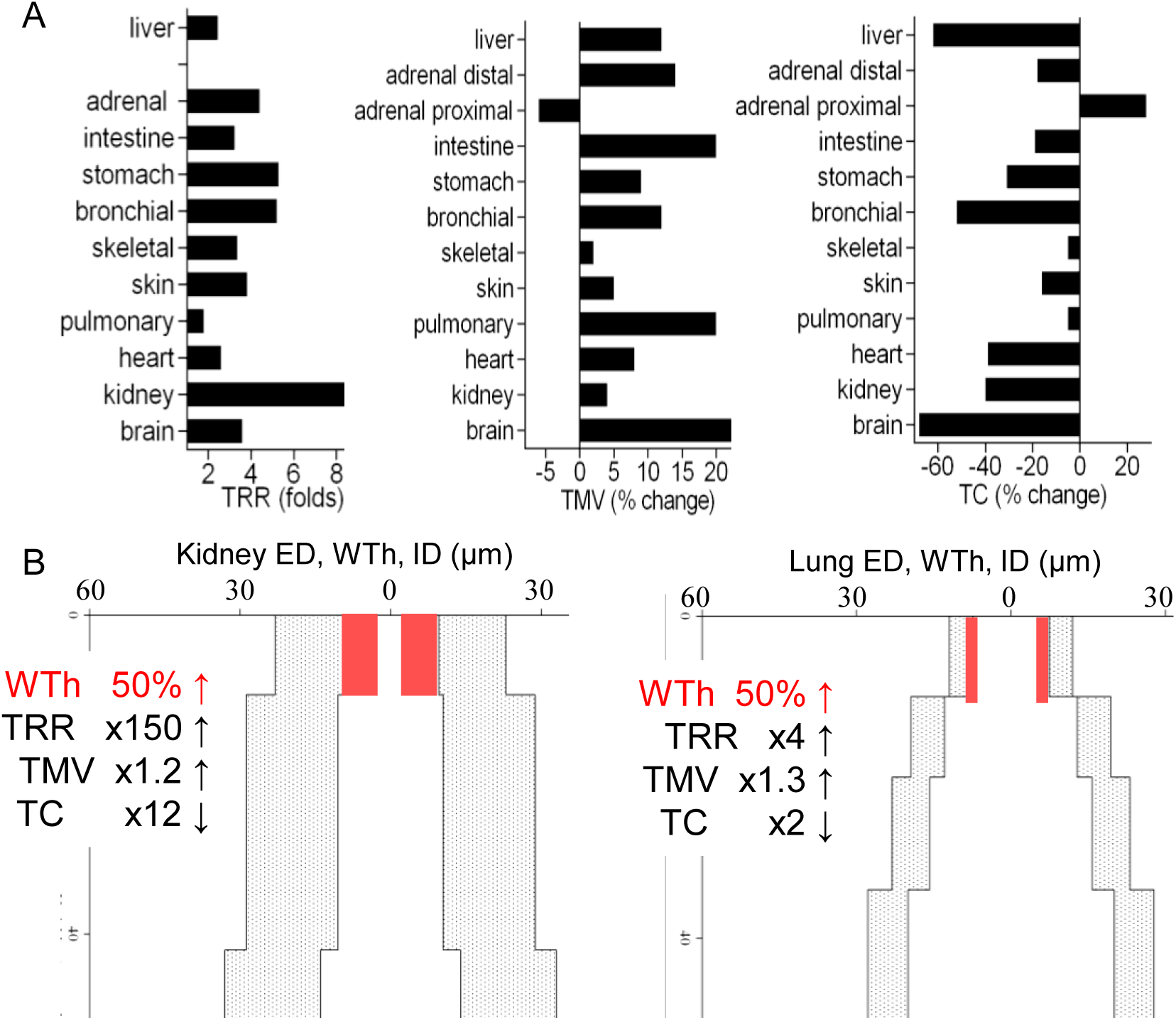
Hemodynamic parameters are enhanced variously in different organs. (A) There was no clear correlation between the values of TRR, TC, TMV. For example, in the brain enhanced TMV corresponded to a large drop in TC but a small increase in TRR. In the kidney a small elevation in TMV corresponded to a small lowering in TC but augmented TRR. (B) An equal 50% increase in wall expansion in kidney and lung arteries would cause marked differences in TRR and TC increases, while TMV would only increase mildly and remain comparable between the two organs. Different dynamics for TRR, TMV, and TC are the result of different initial organ specific dimensions. x - fold increase.

In normal rats TC correlated with the regional BFR (**Fig 10**). In 1K1C rats TC demonstrated maximum decrease in the brain, liver and intestine, and minimal lowering in the skin, pulmonary and skeletal muscle arteries (**Fig 24A**). However, it was difficult to compare our results with pathophysiological studies because regional BFR has been shown as decreased, increased, or stable as well[105],[109],[110]. TMV indicated arterial wall thickening, with maximum of 20-25% in the brain, intestine and pulmonary arteries, and minimum of 2-5% in skeletal muscle, skin and kidney, which were not proportional to TRR and TC (**Fig 24A**). Presumably, basic organ-specific arterial dimensions could have significant impact on hemodynamic consequences of remodeling being the main cause to various organ dynamics of TRR, TC, and TMV (**Fig 24B**). Evidently, hemodynamic correlations of TRR, TC and TMV need further investigation.

An important goal of our work was to determine if random tissue sections are a reliable source of data. Numerous studies of remodeling, exploring random tissue sections have been published in the past three decades, with most in the 1980s or 1990s, so the majority of our references are dated 15-20 years back. Since then *in vitro* myography studies dominate significantly, while morphometric studies on random sections have been infrequent.

Started from the study[51], the post-mortem contraction has been considered a main cause of high variability in arterial morphometry. Perfusion fixation was intended to eliminate post-mortem arterial contraction, especially with preliminary application of a vasodilator[31] or vessel deactivation[111]. The present study indicates that perfusion fixation is not mandatory for morphometry on random tissue sections. In fact, perfused vs non-perfused organs or myograph experiments demonstrated similar stochastic remodeling patterns (**S3 and S4 Table**). Here, we avoided fixation deliberately, performing only immersion fixation in order to have animal (and potentially human) material in the same condition for comparison, since for the latter perfusion is unlikely to be applied. Our data prove that careful consideration of arterial tapering is the most important factor for elaborated morphometry analysis. A similar approach has been widely accepted in atherosclerotic remodeling[93],[57],[112],[113].

In conclusion, we have developed an algorithm to quantify and standardize arterial remodeling analysis. Tissue sampling should be random and representative, according to known recommendations[33],[85]. Following fixation, sections must be stained with periodic acid—Schiff and Masson trichrome. Hematoxylin and eosin staining does not always clearly differentiate muscular and adventitial components, especially when hyalinosis, fibrinoid necrosis, or perivascular inflammatory infiltrates occur[85]. Random measurements of EDs and IDs are performed to obtain about 80-100 measurements of arteries with EDs between ~10-50 μm. Then, variables are arranged in order of increasing ED, divided into 5 μm intervals, and statistical analysis performed for each interval. Finally, the regression equations, complex profiles, remodeling variant, hemodynamic parameters are computed from interval statistics and compared among models or organs. The algorithm does not require additional counting or data gathering, compared to conventional morphometry of arteries on histological sections, and represents a more informative, standardized approach to arterial profiling.

## Material and Methods

Experimental protocols were approved by the Animal Ethics Committee at the University of Ottawa and performed according to the recommendations of the Canadian Council for Animal Care. Analyses in normal rats were performed on 20 male Wistar rats (Charles River, Montreal, Québec, Canada), age 20-25 months, and weighing 600-800 g. Five normal male C57BL6 mice (age 20-30 weeks) were used for comparative morphometry. Goldblatt 1K1C hypertension was induced in 20 male Wistar rats (Charles River Laboratories, Montreal, Québec, Canada), age 20-25 months, weighing 600-800 gm. Under isoflurane anesthesia a silver clip with internal diameter 0.26 mm was placed around the left renal artery and the right kidney was removed. On both sides the manipulation was distal to the adrenal arteries to keep them intact. The control sham-operated group (5 animals) was kept under the same living conditions as the experimental animals. Systolic blood pressure was measured weekly by tail-cuff plethysmography. After 60 days rats were euthanized with an intraperitoneal overdose of sodium pentobarbital. The brain, lung, heart, stomach, liver, small intestine, spleen, kidneys, adrenals, hip skeletal muscle, and tail skin were excised and immersion-fixed in 10% buffered formalin for 24 h, dehydrated and embedded in paraffin. Three fresh lungs from control rats were prepared for special analysis: the entire lobular pulmonary arterial tree was cleaned from parenchyma up to branches of ED 20-30 μm by dissection under a microscope in phosphate-buffered saline. The lung is the only organ which arteries are relatively easy to separate from parenchyma, compare to other organs. Pairs of branches with the same ED but located in proximal and distal segments, 1-2 mm in length were sampled where a branch began, and also embedded in paraffin. Tissue blocks were specifically oriented according to the known anatomical distribution of arteries in order to obtain predominantly cross-circular arterial sections. Histological sections (5 μm thickness) were stained with hematoxylin-eosin and Masson’s trichrome. In sections of three to four blocks from each organ, small arteries and arterioles were traced by visual scanning of the entire section. Only vessels with a long- to short-axis ratio < 1.50 were measured. In this way, the error associated with calculating the diameter by averaging the maximum and minimum diameters was minimized (<3%)[33]. The ED and ID were measured in arteries if the profile had a visibly noninterrupted circular or ellipsoid shape. From those two values the derivatives were calculated as follows: WTh = (ED – ID)/2; WLR = WTh /ID; media cross sectional area (MCSA) = π^*^ (ED^2^/2-ID^2^/2). Microscopy was performed with a Zeiss AX10microscope (Oberkochen, Germany) and images were analyzed by ImagePro Plus software (Media Cybernetics, Bethesda, MD, USA).

Descriptive statistics calculated mean, standard deviation (SD), standard error of mean (SEM), coefficient of variation (CV), and frequency distribution to build histograms and complex profiles. Distribution was assessed by Kolmogorov-Smirnov, D’Agostino & Pearson and Shapiro-Wilk tests. Outliers were identified by the combined robust regression and outlier removal (ROUT) method. Acquired linear regression equations were tested to determine if slopes and intercepts were significantly different (P < 0.05) and the goodness of fit coefficient *r*^2^ was counted for each equation. Nonlinear regression was approximated with exponential growth equations and the goodness of fit coefficient *R*^2^ was counted for each equation. Extra sum-of-squares F-test and Akaike’s Information Criteria (AICc) were used to compare the best-fit values between nonlinear regression equations (P < 0.05). Statistical analyses were performed with GraphPad Prism software (GraphPad Software, La Jolla, CA, USA). Modeling procedures were made with MathCAD version 15.0 (Parametric Technology Corporation, Needham, MS, USA). Three-dimensional graphs were built in OriginPro version 2016 SR0 b9.3.226 (OriginLab Corporation, Northampton, MA, USA).

## Author contributions

Conceived and designed the experiments: AG SA. Performed the experiments: AG SA. Analyzed the data: AG KB PB. Supervision: KB SP. Writing ± review & editing: AG PB KB CK.

## Supporting information captions

**S1 Table.** Normality tests for averaged arterial dimensions in the kidney, heart and pulmonary arteries.

**S1 Fig. Complex profiles revealed different tapering patterns in organs.** Complementary graphs to **Fig 6A**. Axis X – the bidirectional common linear scale for the external diameter (ED), wall thickness (WTh), and internal diameter (ID); axis Y – accumulated frequency of variables (%).

**S2 Fig. Linear regression equations and RR in different organs.** Lines and RR (blue curves) represent the best fit for different organs. Complementary graphs to **Fig 6B**. Points are mean ± SD for 5-μm ED intervals; *r*^2^ - goodness of fit coefficients. Corresponding equations are in **S2 Table.**

**S2 Table. Linear regression equations from mean ± SD for 5 μm ED intervals.**

**S3 Fig. Comparison of linear regressions for ID and WTh between different organs.** Equations for lungs, bronchi, adrenal glands, stomach and skeletal muscles were very distinctive (P<0.0001). Brain and intestine (*) shared similar equations (P>0.41 for ID and WTh). Heart and spleen (**) were also close (P>0.43 for ID and >0.83 for WTh). Equations for kidney, liver and skin (***) were similar (P>0.77 for ID and >0.87 for WTh).

**S4 Fig. Histograms of terminal arteries in the kidney, heart and brain.** The significant irregularity and asymmetry for dimensions used to calculate statistics in **Table 2**. Data did not pass conventional statistical tests for normality (negative, P<0.001).

**S3 Table. Numerical classification applied to arteries studied via in vitro myography**. 1K1C, one kidney-one clip; 2K1C, two kidney-one clip; Ang, angiotensin; DM2, diabetes mellitus type 2; EHT, essential hypertension; eNOS, endothelial nitric oxide synthase; GI, growth index; L-NAME, Nitro-L-arginine methyl ester; op/+, osteopetrotic heterozygous; PM, pressure myograph; RI, remodeling index; RVH, renovascular hypertension; SA, statistical artifact; SHR, spontaneously hypertensive rats; SHRSP, spontaneously hypertensive rats stroke prone; SOD, superoxide dismutase; WM, wire myograph.

**S4 Table. Linear regression equations of terminal arteries derived from mean ± SD for 5-μm ED intervals**. All equations for hypertensive rats were significantly different from equations for control rats (*r*^2^ = 0.99; P < 0.0001).

**S5 Fig. Control vs hypertensive complex profiles in organs.** Control (green) and hypertensive (red) complex profiles were superimposed. Remodeling patterns are not recognizable. Complementary graphs to **Fig 18A**.

**S6 Fig. Linear regression lines displaced similarly in most organs**. The ID slopes decreased (solid arrows), and the WTh slopes increased (dashed arrows). Adrenal arteries demonstrated opposite directions: the increased ID slopes and decreased WTh slopes. Complementary graphs to **Fig 18B**.

**S7 Fig. The remodeling of renal arteries has been simulated for variants 2-4c, 2-6a, all 3-4, 3-5a, and 3-6a**. Displacement of linear regression lines up or down for any parameter was similar for many remodeling variants. Complementary graphs to **Fig 19**.

**S8 Fig. 3D-modeling simulation for the brain**. No variants were congruent. Complementary graphs to **Figure 20**.

**S9 Fig. 3D-modeling simulation for the distal arterial segments in the adrenals**. No variants, except 2-4c, were congruent. Complementary graphs to **Fig 22A**.

**S10 Fig. 3D-modeling simulation for the proximal arterial segments in the adrenals**. If the variant 1-6aI appeared, congruence to the variant 2-6a must also be present because data sets overlap. Complementary graphs to **Fig 22B**.

**S11 Fig. 3D-modeling simulation for the kidney.** If the variant 3-4cIII occurred, simultaneous congruence to 2-4c and 3-4cII (but not 3-4cI) must be present because data sets overlap. Complementary graphs to **Fig 22C**.

**S5 Table. Averaged arterial dimensions in rats with aortic coarctation and 1K1C hypertension** (modified from[102]). NC, numerical classification; SA – statistical artifact; ↑ indicates increase in dimensions vs sham values, *P<0.05, ** P<0.01.

**S6 Table. Averaged arterial dimensions in rats with aortic coarctation and 2K1C hypertension** (modified from[17]). 2K1C, two-kidney, one clip; NC, numerical classification; SA – statistical artifact; ↑ ↓ indicates increase or decrease in dimensions vs sham values, *P<0.05, ** P<0.01, *** P<0.001

**S12 Fig. Cremaster arteries in normotensive rats with aortic coarctation have no remodeling**. The control and experimental equations were not different (the variant 2-5b; P > 0.6-0.7). Calculated from[102]. Complementary graphs to **Fig 23A**.

**S13 Fig. Renal interlobular arteries in normotensive rats with aortic coarctation have no remodeling**. The control and experimental equations were not different (the variant 2-5b; P > 0.8). Calculated from[17]. Complementary graphs to **Fig 23A**.

**S14 Fig. Cremaster arteries in hypertensive 1K1C rats developed the variant 2-4c**. Calculated from[102]. Complementary graphs to **Fig 23B**.

**S15 Fig. Renal arteries in 2K1C rats were congruent to the variant 2-4c**. Calculated from[17]. Complementary graphs to **Fig 23C**.

**S16 Fig. Renal arteries in SHR showed congruence to the variant 2-4c**. Calculated from[39]. Complementary graphs to **Fig 23D**.

**S17 Fig. Pulmonary arteries in a case of pulmonary hypertension due to congenital mitral stenosis developed the variant 1-5c**. Calculated from[55]. Complementary graphs to **Fig 23E**.

**S18 Fig. Recalculated data from human subcutaneous arteries** [104]. The graph length vs area has been approximated with the linear regression, resulting in curved lines. Complementary graphs to **Fig 23F**.

**S19 Video. 3-D modeling for the brain**. The video demonstrates the variant 2-4c as congruent.

**S20 Video. 3-D modeling for the brain**. The video demonstrates the variant 3-4aIII as an example of a non-congruent variant.

